# Maternal Transfer of Environmentally Relevant Polybrominated Diphenyl Ethers (PBDEs) Produces a Diabetic Phenotype and Disrupts Glucoregulatory Hormones and Hepatic Endocannabinoids in Adult Mouse Female Offspring

**DOI:** 10.1101/2020.08.31.275008

**Authors:** Elena V. Kozlova, Bhuvaneswari D. Chinthirla, Pedro A. Pérez, Nicholas V. DiPatrizio, Donovan A. Argueta, Allison L. Phillips, Heather M. Stapleton, Gwendolyn M. González, Julia M. Krum, Valeria Carrillo, Anthony E. Bishay, Karthik R. Basappa, Margarita C. Currás-Collazo

## Abstract

Polybrominated diphenyl ethers (PBDEs) are brominated flame retardant chemicals and environmental contaminants with endocrine-disrupting properties that are associated with diabetes and metabolic syndrome in humans. However, their diabetogenic actions are not completely characterized or understood. In this study, we investigated the effects of DE-71, a commercial penta-mixture of PBDEs, on glucose regulatory parameters in a perinatal exposure model using female C57Bl/6 mice. Results from *in vivo* glucose and insulin tolerance tests and *ex vivo* analyses showed that DE-71 produced fasting hyperglycemia, glucose intolerance, reduced sensitivity and delayed glucose clearance after insulin challenge, and exaggerated hepatic endocannabinoid tone in F1 offspring exposed to 0.1 mg/kg DE-71 relative to control. DE-71 effects on F0 dams were more limited indicating that indirect exposure to developing offspring is more detrimental. Other *ex vivo* glycemic correlates occur more generally in exposed F0 and F1, i.e., reduced plasma insulin and altered glucoregulatory endocrines, exaggerated sympathoadrenal activity, decreased thermogenic brown adipose tissue mass and reduced hepatic glutamate dehydrogenase enzymatic activity. Hepatic PBDE congener analysis indicated maternal transfer of BDE-28 and −153 to F1 at a collective level of 200 ng/g lipid, in range with maximum values detected in serum of human females. Given the persistent diabetogenic phenotype, especially pronounced in female offspring after developmental exposure to environmentally relevant levels of DE-71, additional animal studies should be conducted that further characterize PBDE-induced diabetic pathophysiology and identify critical developmental time windows of susceptibility. Longitudinal human studies should also be conducted to determine the risk of long-lasting metabolic consequences after maternal transfer of PBDEs during early-life development.

## Introduction

Polybrominated diphenyl ethers (PBDEs) are a class of anthropogenic persistent organic pollutants (POPs) that have been added to polymers and textiles since the 1970s to reduce the flammability of commercial products such as furniture, building materials and electronics^1^. PBDEs have been marketed as industrial mixtures, varying in percent composition of congeners based on the number of bromine substitutions on the phenyl rings; forming 209 theoretical congeners. These chemicals are lipophilic, highly resistant to degradation and are easily released from products into the indoor environment and bioaccumulate up the human food chain^2,3^. Over a 30-year period, PBDE levels have increased exponentially in adult human tissues including blood, adipose^4^, organs^5^, breast milk and fetal^6^ and child tissues sampled worldwide^7^. Despite legislative action by the European Union and a voluntary phase-out of production in the US starting in 2005, which led to a decrease in environmental concentrations, PBDE contamination remains an ongoing problem since products containing PBDEs are still in circulation and re-entering the anthroposphere from electronic waste sites^8^ and inadvertent recycling^9^. PBDE levels in various sample types worldwide, including the breastmilk and sera of U.S. women and toddlers are still being detected^10,11,12,13^. Moreover, modelling studies predict that penta-, octa- and deca-brominated BDEs will continue to be emitted from in-use and waste stocks until 2050^9^. Given that PBDEs are still accumulating in humans, it is important to determine the long-term health consequences of chronic exposures.

Human and animal studies have associated developmental PBDE exposure with endocrine disruption, especially impaired thyroid hormone homeostasis^14,15^, neurotoxicity^15,16,17^ and lower birth weight and length^18^. Stressful physiological conditions during development can cause a predisposition to chronic disease later in life^19^. Such suboptimal conditions during perinatal life include exposure to PBDEs due to mobilization from maternal sources through cord blood *in utero^20^* and breast milk during lactation^11,21^. After birth, toddlers continue to be exposed to environmental PBDEs through household dust and diet. These factors, as well as immature detoxification in the liver, contribute to a 3 to 9-fold higher PBDE body burden in infants and children as compared to adults^22,23^. Thus, developmental exposure to highly penetrant and active environmental xenobiotics such as PBDEs is of high concern due to potential significant health risks posed during developmental time points sensitive to biological reprogramming.

Type 2 diabetes (T2D) has shown a dramatic rise in incidence in recent years. It is estimated to become the greatest epidemic of the 21^st^ century; with a predicted increase to 592 million affected by 2035^24^. Epidemiological studies suggest that the escalating production and environmental presence of a number of metabolic disrupting chemicals (MDC) over the past four decades may contribute to the pathogenesis of metabolic diseases^25^. Mounting evidence has implicated brominated POPs in the pathogenesis of T2D and metabolic syndrome (MetS)^26,27,28, 29,30,31,32,33,34^. In particular, diabetes and/or MetS are positively associated with high body burdens of individual PBDE congeners including: BDE-28, −153, and −47, one of the most abundant PBDE congeners detected in the environment and in human tissues^29,34,35,36^. Additionally, exposure to BDE-28, −47, −99, −153, −154 and −183 increases the risk of gestational diabetes mellitus (GDM) in healthy US women^37,38,39^, a physiologically demanding condition that may increase the risk of MetS in adult offspring^40^. All of the PBDE congeners that have been associated with T2D are found in DE-71, a commercial mixture of PBDEs with high environmental and human relevance^41^. DE-71 congeners are shown to accumulate in liver and adipose tissue^17^, which are critical for glucose homeostasis.

Given that developing organisms are more susceptible to potential metabolic disruptors such as PBDEs, it is important to test whether they predispose offspring to diabetes in adulthood. To answer this question, we employed a mouse model of chronic, low-dose maternal transfer of environmentally relevant PBDE congeners to characterize the adult consequences of exposure during critical perinatal developmental windows. We tested the hypotheses that exposure to DE-71 produces a diabetogenic phenotype and that DE-71 effects are more pronounced in developmentally exposed female offspring versus their adult-exposed mothers. We focus on female mice, since impaired glucose tolerance is more common in diabetic women who also have an increased risk of death compared to non-diabetics^42^. Our results indicate that exposure to DE-71, especially when administered perinatally, alters clinically relevant diabetic biomarkers, namely, fasting blood glucose, glucose tolerance, insulin sensitivity, plasma levels of glucoregulatory hormones as well as liver endocannabinoid tone, an emerging biomarker of energy balance. These findings raise concern for the health of progeny of directly exposed mothers, especially since the diabetogenic effects of DE-71 involve multiple organ system biomarkers and persist into adulthood.

## Results

C57Bl/6 mouse dams were exposed to DE-71 and later investigated along with their offspring **(Fig. 1**).

**Figure 1.**
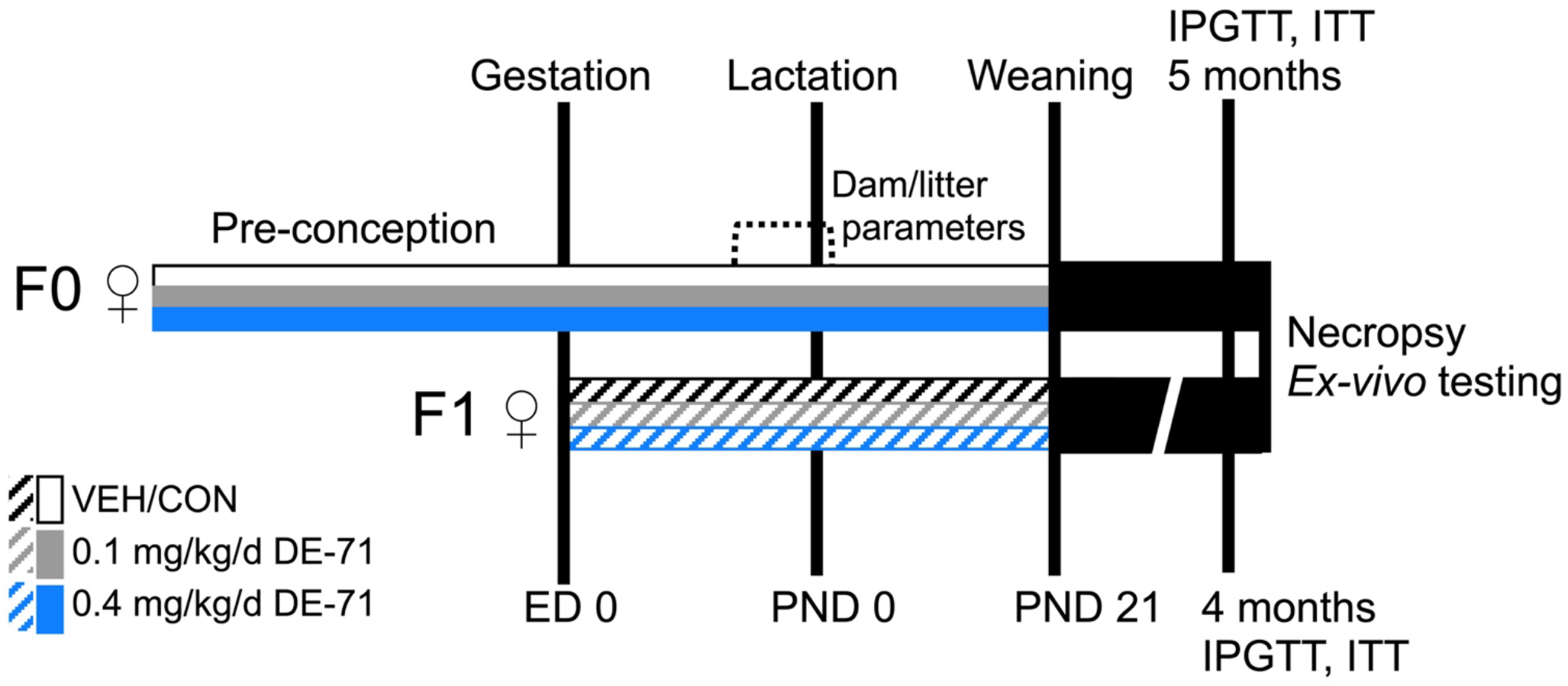
Dosing and testing paradigm used for perinatal and adult exposure to DE-71. Direct exposure to DE-71, represented by solid shading in adult dams (F0♀), began ~4 weeks pre-conception and continued daily for ~3 weeks of gestation and 3 weeks of lactation until pup weaning at PND 21. Indirect exposure to DE-71, represented by hatched shading, in female offspring (F1♀) occurred perinatally (ED 0 to GD 18) and postnatally via lactation through weaning (PND0-PND21) for ~39 d. Maternal and litter parameters were taken during GD 15 to PND 0. Using this regimen, exposure duration was 70-80 d for F0 and 39 d for F1. Metabolic endpoints (fasting glycemia, IPGTT, ITT) were examined in adulthood, at 1-2 weeks post-weaning for dams (at ~5 months old) and ~4 months for offspring. At necropsy, body and organ weights were recorded and blood and organ tissues collected for further analysis: GC/ECNIMS BDE congener determination, plasma ELISA, UPLC/MS/MS detection of endocannabinoids, epinephrine assay, liver enzymatic activity. GD, gestational day; PND, postnatal day; ED, embryonic day

### Dam gestational parameters and litter outcomes

DE-71 exposure did not significantly affect maternal food intake, weight gain, litter size or secondary sex ratios (see **Supplementary Table S1 online**).

### Congener analysis in livers of F0 and F1 females

Notable differences were observed in hepatic BDE congener accumulation after direct vs indirect exposure to DE-71 in F0 and F1 females, respectively. The sum of detected BDE congeners (∑PBDEs, geometric mean+SD) in dams was approximately 15-fold greater relative to offspring exposed to 0.1 mg/kg DE-71 (**Fig. 2a**; *P*<0.05). An effect of increasing dose was seen only in F1. Individual congener analysis indicates only tri- (BDE-28/33) and hexa-brominated (BDE-153) congeners were detected in offspring liver. In contrast, more congeners were detected in dams: tri- (BDE-28/33), tetra- (BDE-47, −66), penta- (BDE-85, −99, −100), hexa- (BDE-153, −154) and hepta-brominated congeners (BDE-183) (**Fig. 2c, d**). Figure 2b shows that, of the total congeners measured, the predominant ones in the dam liver were (% in 0.1 - % in 0.4 mg/kg): BDE-99 (24-43%) and BDE-153 (21-53%). The overall congener pattern observed in dam liver was similar to that in DE-71^17,41^ except for the greater content of BDE-153 and the lower content of BDE-47 (6-14%) and BDE-100 (8.5-14%). In combination, these four congeners accounted for 92% (0.1 mg/kg) and 91.5% (0.4 mg/kg) of total BDEs in F0 liver. In contrast, BDE-153 accounted for 97% and 94% of total PBDEs detected in F1 exposed to 0.1 mg/kg and 0.4 mg/kg doses, respectively. The exaggerated penetration of BDE-153 (4 to 10-fold greater in F0 and 17 to 18-fold greater in F1 than the congener profile of DE-71) is consistent with previous reports^43^. For complete congener profiles see **Supplementary Table S2 online**.

**Figure 2.**
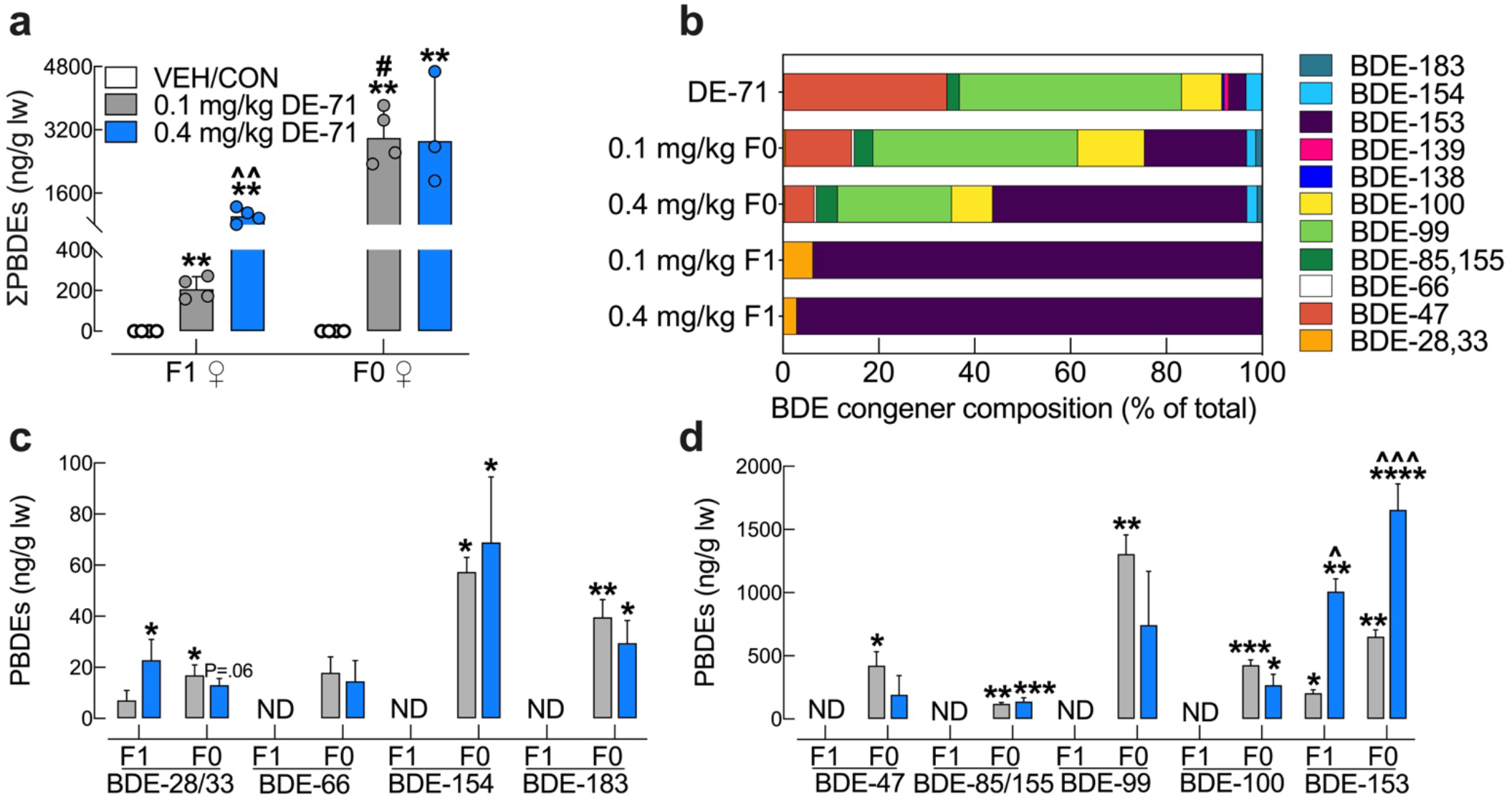
Hepatic BDE congener analysis of DE-71-exposed mouse dams and their adult female offspring. (**a**) The ng/g lipid wt sum concentrations of PBDE congeners detected (∑PBDEs; geometric mean±geometric SD) comprised of BDE 28/33, BDE-153 for F1 and BDE-28/33, BDE-47, BDE-66, BDE-85/155, BDE-99, BDE-100, BDE-153, BDE-154, BDE-183 for F0. DE-71 exposure produced significant accumulation of PBDEs in liver, being greater in directly exposed F0 versus indirectly exposed F1 female mice. Dose-dependency was only seen in F1. Values for VEH/CON are not shown since they were below the method detection limit (MDL). (**b**) BDE composition (percent total) in the lot of DE-71 used and in livers of F1 and F0. The multi-congener profile in F0 was similar to that of DE-71, whereas that of F1 was restricted to BDE-28/33 and BDE-153. Co-elution of BDE −28 and −33 as well as BDE-85 and −155 prevented differentiation during analysis. (**c, d**) Absolute concentrations of congeners found in F1 and F0 liver. Predominant congeners in F0 were BDE-47, −99, −100, and −153. Predominant congeners in F1 were BDE-28/33 and −153. Only BDE-153 showed a rise in content in F0 and F1 mice exposed to 0.4 mg/kg relative to 0.1 mg/kg. For statistical purposes, values below the MDL were substituted with randomly generated values between 0 and MDL/2 and designated as not detected (ND). Bars and error bars reflect mean±s.e.m. *indicates significantly different from VEH/CON (\**P*<.05, \*\**P*<.01, \*\*\**P*<.001, \*\*\*\**P*<.0001); ^indicates significantly different from the corresponding 0.1 mg/kg group (^*P*<.05, ^^*P*<.01, ^^^*P*<.001). #significant difference across F0 and F1. Dunnet’s T3 or Tukey’s *post-hoc* tests were used. n=3-4 replicates/group, analyzed in triplicate. F1, female offspring; F0, dams; ND, not detected

### Chronic low dose DE-71 exposure has minimal effects on body and selected organ weights

Body weights of female offspring perinatally exposed to 0.1 mg/kg were significantly lower by approximately 7% relative to VEH/CON (see **Supplementary Table S3 online**). Absolute liver weight was greater in 0.4 mg/kg offspring relative to VEH/CON (9%) and 0.1 mg/kg (13%). The absolute and relative weights of pancreas and spleen of the F1 females were similar across groups. The only difference seen in dams was the 9% greater relative liver weight of 0.1 mg/kg dams compared to VEH/CON. Fasting body weights taken from a subset of mice used for IPGTT and ITT were not different across groups and, therefore, the diabetogenic phenotype of DE-71-exposed F1 mice is not due to obesity.

### DE-71 produces fasting hyperglycemia in F1 but not F0 females

One indication of pre-diabetes is abnormally high fasting blood glucose (FBG) concentration^44,45^. We examined FBG after 9 and 11 h fasting using glycemia values from basal time points obtained in IPGTT and ITT experiments. For the ITT, we used a 9 h fast time and a corresponding low insulin dose (0.5 U/kg). In female offspring, 0.1 mg/kg DE-71 significantly elevated FBG after a 9 h fast and 0.4 mg/kg DE-71 elevated FBG after an 11 h fast relative to VEH/CON (**Fig. 3a**). Therefore, hyperglycemia was present in exposed F1 at both fast times, albeit the effective DE-71 dose was different depending on fasting duration. In contrast, DE-71 exposure did not significantly affect FBG in F0 females, regardless of the fasting condition (**Fig. 3b**). These results suggest that perinatal exposure to DE-71 produces fasting hyperglycemia, a condition that may be due to dysregulated endocrine parameters of glucose metabolism^46^.

**Figure 3.**
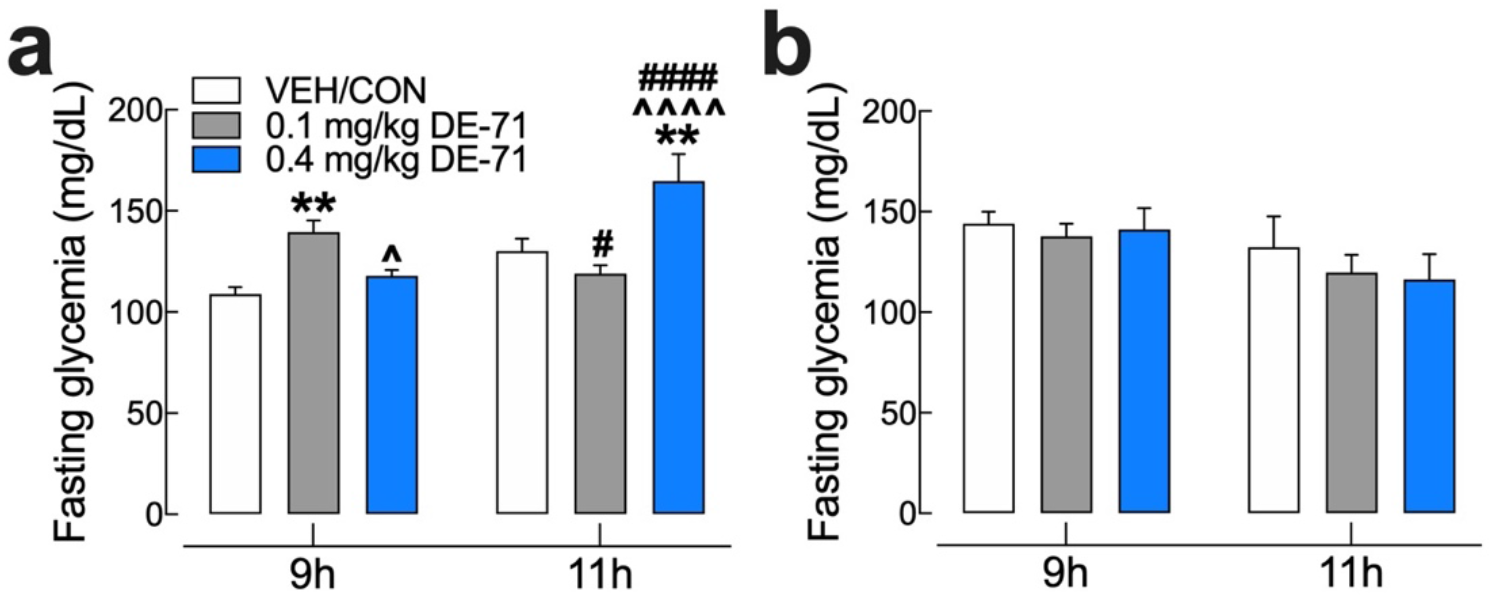
DE-71 exposure produces elevated fasting blood glucose (FBG) in perinatally exposed female offspring but not their mothers. FBG was measured after a 9 or 11h fast in female offspring (**a**) and dams (**b**). *indicates significantly different from VEH/CON (\*\**P*<.01); ^indicates significantly different from 0.1 mg/kg DE-71 (^*P*<.05, ^^^^*P*<.0001). ^#^indicates significant difference between 9 and 11 h for corresponding exposure group (#*P<.05*; ####*P*<.0001). Sidak’s *post hoc* test was used. Bars and error bars represent mean±s.e.m. n=7-12/group. F1, female offspring; F0, dams. FBG, fasting blood glucose.

### DE-71 exposure impairs glucose tolerance after developmental or adult exposure

To investigate the effects of DE-71 on glucose tolerance, glycemia was measured during IPGTT. Blood glucose levels rose rapidly and peaked within 15 min of glucose challenge in the VEH/CON and the 0.4 mg/kg DE-71 groups. In contrast, the corresponding peak for the 0.1 mg/kg DE-71 group occurred later, at 30 min (**Fig. 4a**). Relative to VEH/CON, the glycemia was exaggerated in exposed F1 at 30 and 60 min (0.1 mg/kg) and at 15 min post injection (0.4 mg/kg) indicating glucose intolerance, with an especially pronounced magnitude in the 0.1 mg/kg DE-71 group (**Fig. 4a**). Plasma glucose showed a gradual return to baseline at 60 min post injection in VEH/CON. In contrast, for both exposed F1 groups, the corresponding time was 120 min, or 1 h longer (**Fig. 4a**)‥ The differences in magnitude and duration of glycemia are incorporated in the area under the glucose curve, AUC_IPGTTglucose_, which is abnormally large in F1 females exposed to either dose (**Fig. 4b**). The latency to maximum glycemia was not significantly different across groups (**Fig. 4c**). Because FBG after an 11 h fast is elevated in F1 exposed to 0.4 mg/kg, glycemia, values are also expressed as percent baseline (**Fig. 4g)**. In this case, similar results were found to those expressed using absolute glycemia values but only the 0.1 mg/kg dose group shows significantly greater AUC_IPGTTglucose_ (**Fig. 4h**).

**Figure 4.**
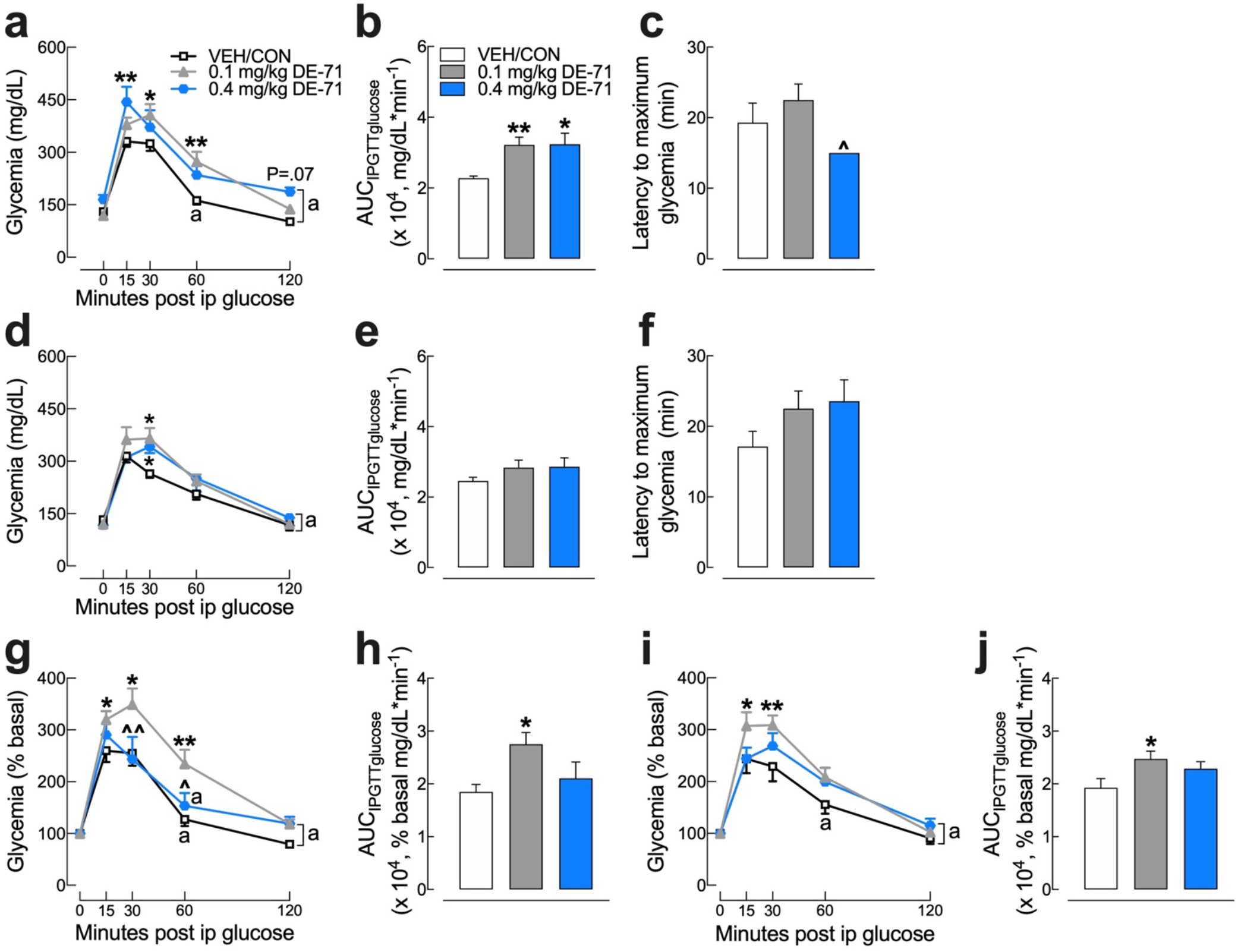
DE-71 exposure produces greater glucose intolerance after perinatal exposure compared to adult exposure. Mice were fasted for 11 h ON and tail blood was sampled for glucose before (t=0 min) and after (t=15, 30, 60 and 120 min) i.p. injection of 2.0 g/kg glucose. Absolute blood glucose concentrations taken during IPGTT of F1 (**a**) and F0 (**d**). Mean values for the integrated area under the IPGTT glucose curve (AUC_IPGTTglucose_) for F1 (**b**) and F0 (**e**). Blood glucose values taken during IPGTT are plotted vs time as a percent of basal glucose for F1 (**g**) and F0 (**i**). Mean values for integrated area under the IPGTT glucose percent basal curve (AUC_IPGTTglucose_) for F1 (**h**) and F0 (**j**). Latency to maximum glycemia for F1 (**c**) and F0 (**f**). *indicates significantly different from VEH/CON (\**P*<.05, \*\**P*<.01); ^indicates significantly different from 0.1 mg/kg DE-71 (^*P*<.05, ^^*P*<.01,). Tukey’s or Dunnet’s *post hoc* tests were used. The symbol “a” indicates the time points at which glycemia is not different from basal in the corresponding treatment group. Glycemia at all other time points differs from basal. Bars and error bars represent mean±s.e.m. n=7-12/group. F1, female offspring; F0, dams

Exposed F0 also showed significantly greater glycemia relative to VEH/CON but the difference was moderate and this occurred at 30 min post-injection (**Fig. 4d).** Glycemia levels (expressed as percent of basal) return to normal by 60 min in VEH/CON but not until 120 min in F0 exposed to either dose group **(Fig. 4i**). In addition, F0 exposed to 0.1 mg/kg show a significantly greater AUC_IPGTTglucose_ relative to VEH/CON (**Fig. 4j**). To test the hypothesis that DE-71-provoked glucose intolerance is exaggerated in F1 relative to F0, we compared percent basal AUC_IPGTTglucose_ for the 0.1 mg/kg exposure groups and found no significant differences (*P*=.17). These results suggest that exposure to DE-71 causes glucose intolerance after either developmental or adult exposure.

### DE-71 exposure produces an abnormal glycemic response to insulin in F1 females

Next, we examined the glycemia response to exogenous insulin during ITT experiments. Mean glycemia values over the 120 min period following insulin injection are shown in the insulin tolerance curve (**Fig. 5a).** F1 exposed to 0.1 mg/kg DE-71 display less reduction in glycemia as compared to VEH/CON at several time points post injection (*t*=15, 30 min). However, this was confounded by the elevated FBG for F1 exposed to 0.1 mg/kg (**Fig. 3a**). Therefore, a more valid group comparison is shown when expressing glycemia as a percent of baseline (**Fig. 5i**). In this case, exposed F1 displayed a deeper insulin curve with a longer recovery time after insulin injection. This is represented as a greater mean latency to reach the minimum insulin-induced hypoglycemia in exposed F1 groups, 72.5 min (0.1 mg/kg) and 70 min (0.4 mg/kg), relative to VEH/CON, i.e., 37.5 min (**Fig. 5d**), possibly indicating delayed glucose clearance/utilization in response to insulin (**Fig. 5d**).

**Figure 5.**
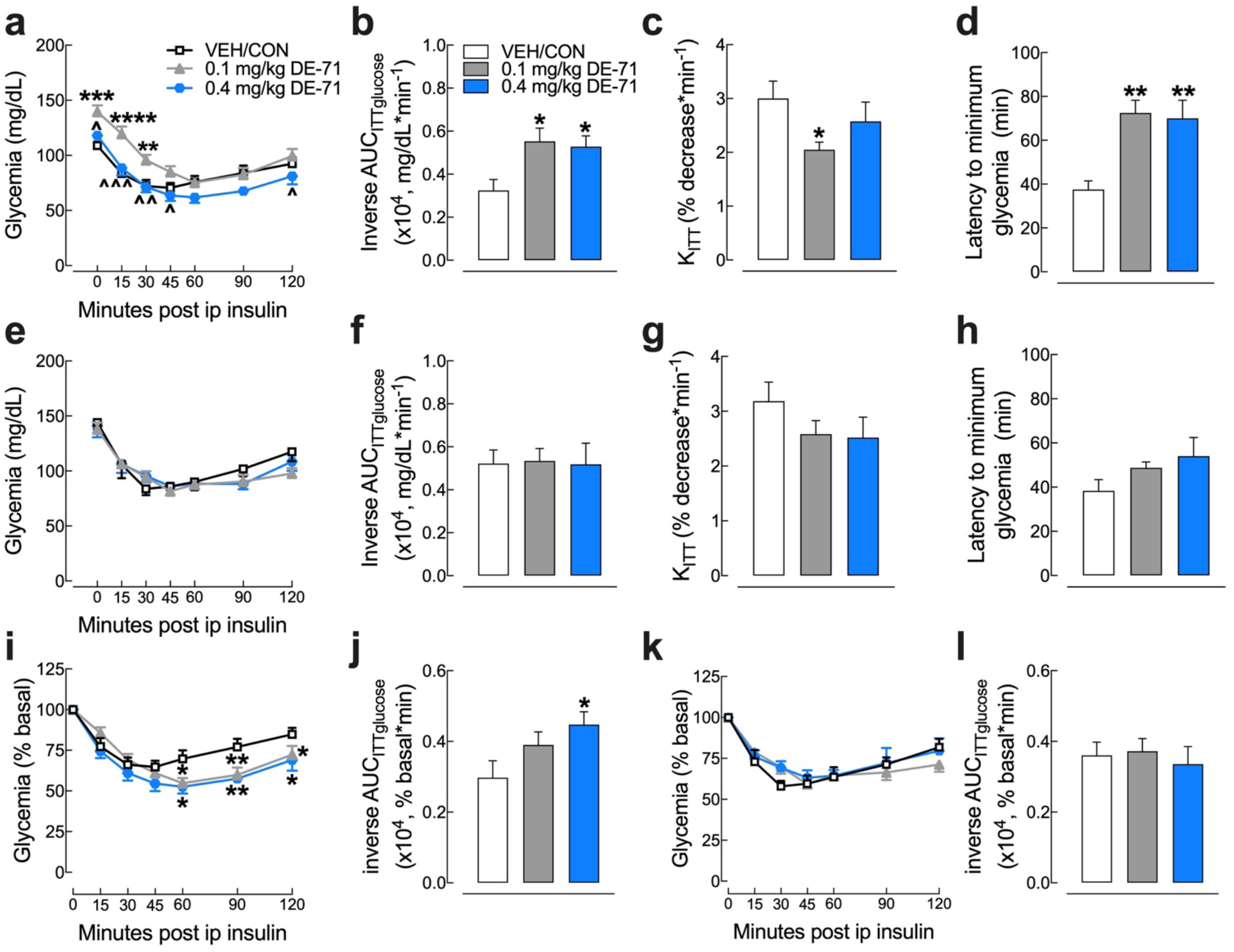
DE-71 exposure causes less glycemia reduction and delayed glucose clearance after insulin challenge in F1 but not F0 female mice. Absolute blood glucose concentrations were recorded before and at *t*=15, 30, 45, 60, 90 and 120 min post-injection with 0.5 U/kg insulin for female female offspring (F1) (**a**) and their mothers (F0) (**e**). Glycemia was analyzed by inverse integrated area under the ITT glucose curve (AUC_ITTglucose_) for F1 (**b**) and F0 (**f**). Rate constant for glucose reduction (K_ITT_) was calculated over the initial slope of ITT glucose response curve from 0-15 min post-injection for F1 (**c**) and F0 (**g**). Latency to minimum blood glucose measured over the two hour time course of ITT glucose response curve for F1 (**d**) and F0 (**h**). Glucose values taken during ITT are plotted vs time as a percent of the individual baseline for F1 (**i**) and F0 (**k**). The inverse integrated area (AUC) under the percent basal glucose curve (AUC_ITTglucose_) for F1 (**j**) and F0 (**l**). *indicates significantly different from VEH/CON (\**P*<.05; \*\**P*<.01, \*\*\**P*<.001, \*\*\*\**P*<.0001). ^indicates significantly different from 0.1 mg/kg DE-71 (^*P*<0.05, ^^*P*<.01, ^^^*P*<.001). Glycemia at all time points differs from basal for corresponding group. Dunnet’s and Tukey’s *post-hoc* tests were used. All values represent mean±s.e.m. n=8-12/group. F1, female offspring; F0, dams

The inverse AUC_ITTglucose_ using percent baseline glycemia showed a significant increase for the 0.4 mg/kg exposed F1 group relative to VEH/CON (**Fig. 5j**). Because glycemia responses over the 120 min observation period is due to complex actions of insulin (insulin signaling at its targets (sensitivity), half-life, and glucose utilization/clearance) we measured early effects of insulin (sensitivity), represented using K_ITTinsulin_ measured over the first 15 min post-injection (**Fig. 5c**). This metric showed a significant decrease (32%) in blood glucose reduction rate for 0.1 mg/kg F1 (*P*=.04) although not 0.4 mg/kg F1 (14%) relative to VEH/CON, suggesting that DE-71 exposure at 0.1 mg/kg produces significant insulin insensitivity.

In contrast to that of F1, the ITT curve for exposed F0 looks normal (**Fig. 5e,k**). No group differences for F0 were observed for K_ITTinsulin_ (**Fig. 5g**). The mean decrease in K_ITTinsulin_ values relative to VEH/CON was 19 and 21% greater for F0 exposed to 0.1 and 0.4 mg/kg DE-71, respectively. In addition, exposed F0 showed normal latency to reach minimum insulin-induced hypoglycemia relative to VEH/CON (**Fig. 5h**). Accordingly, no group differences were seen for inverse AUC_ITTglucose_ (**Fig. f,l**). Taken together, these results indicate reduced insulin sensitivity (0.1 mg/kg) and delayed glucose clearance/utilization (0.1 and 0.4 mg/kg) in response to exogenous insulin injection in exposed F1 but not F0. These results also suggest that developmental, but not adult exposure to DE-71, produces insulin insensitivity characterized by a delay in reaching a peak response to and delayed recovery from insulin challenge, suggesting a diabetogenic phenotype.

### Endocrine-disrupting effects of DE-71 exposure on glucoregulatory hormones

Having observed disruptions in glucose homeostasis we measured plasma hormones involved in carbohydrate regulation using EIA. Plasma insulin levels were significantly lower in F1 exposed to 0.1 mg/kg DE-71 and trending lower in those exposed to 0.4 mg/kg mice relative to VEH/CON (**Fig. 6a**). Exposed F1 females also showed lower plasma glucagon at 0.4 mg/kg but showed no changes in GLP-1 relative to VEH/CON. In exposed F0, mean plasma insulin was also downregulated in 0.1 mg/kg dose group and mean plasma glucagon was trending higher in 0.4 mg/kg dose only. F0 also showed significantly upregulated plasma GLP-1 at 0.1 mg/kg DE-71. Hence, the 0.1 mg/kg F1 group showed the most prominent diabetogenic phenotype and also the most downregulated levels of insulin. At 0.4 mg/kg, DE-71 produces a less pronounced glucose intolerance and an apparent reduction in insulin and reduced glucagon.

**Figure 6.**
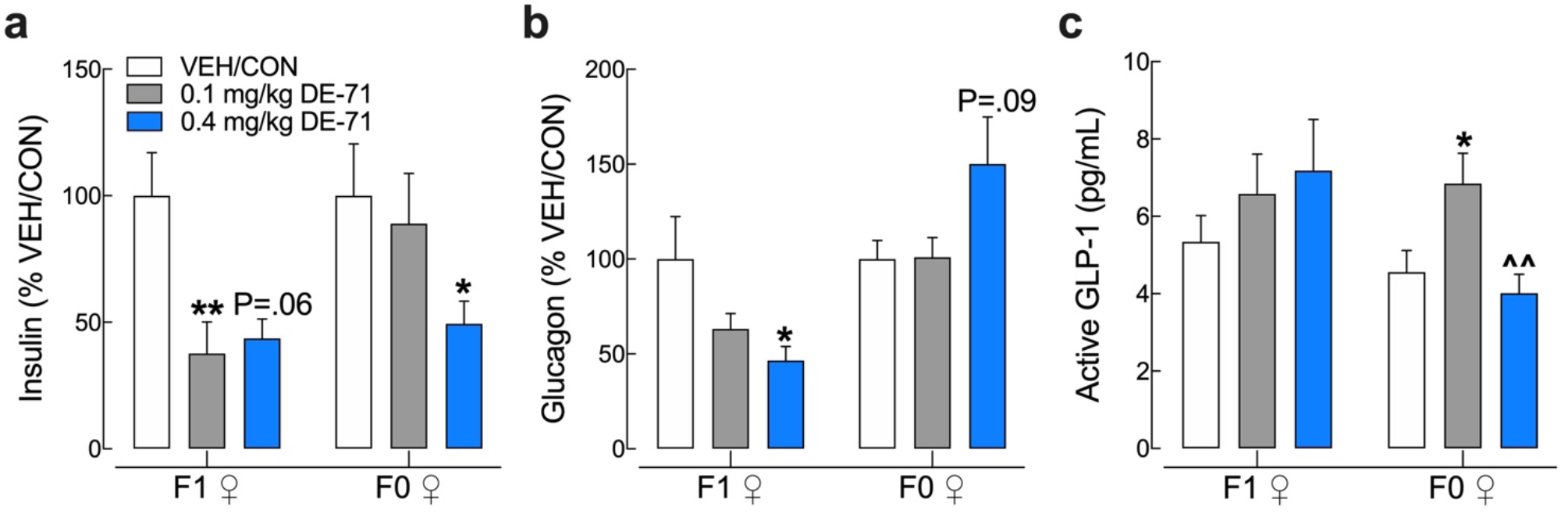
Endocrine-disrupting effects of DE-71 on glucose regulatory hormones in F0 and F1 female mice. Blood collected at sacrifice was assayed for plasma levels of insulin (**a**), glucagon (**b**), and GLP-1 (**c**) in offspring and dams using specific EIA. F1 exposed to DE-71 displayed reduced insulin (1.0 mg/kg) and an apparent reduction in insulin and reduced glucagon (0.4 mg/kg). F0 exposed to DE-71 displayed reduced plasma insulin and apparent rise in glucagon (0.4 mg/kg). *indicate significantly different from corresponding VEH/CON (\**P*<.05, \*\**P*<.01). ^indicates significantly different from 0.1 mg/kg DE-71 (^^*P*<.01). Bars and error bars represent mean±s.e.m. Insulin and glucagon levels are expressed as a percent of VEH/CON. Dunnet’s T3 Dunn’s and Tukey’s *post hoc* tests were used. n=12-24/group for insulin, n=6-8/group for glucagon and n=8-14/group for GLP-1. EIA, enzyme-linked immunoassay; F1, female offspring; F0, dams

### Upregulated adrenal epinephrine content and reduced BAT after DE-71 exposure

Due to the important role of epinephrine in glucose and lipid homeostasis we examined if adrenal content was altered in glucose dysregulated mice exposed to DE-71. Control epinephrine levels were similar to those reported previously for adrenal gland in male mice^47^. Exposure to DE-71 significantly elevated adrenal epinephrine in both dams and female offspring, especially at 0.1 mg/kg dose (**Fig. 7a**). Adrenal weights were no different across experimental groups (data not shown). Brown adipose tissue (BAT) activity increases energy expenditure and has been inversely associated with diabetes and fasting glucose level^48^. When normalized to body weight mean intrascapular BAT mass was significantly decreased by 19% in 0.1 mg/kg exposed F1 relative to VEH/CON (**Fig. 7b**). There were no significant differences due to DE-71 exposure in F0.

**Figure 7.**
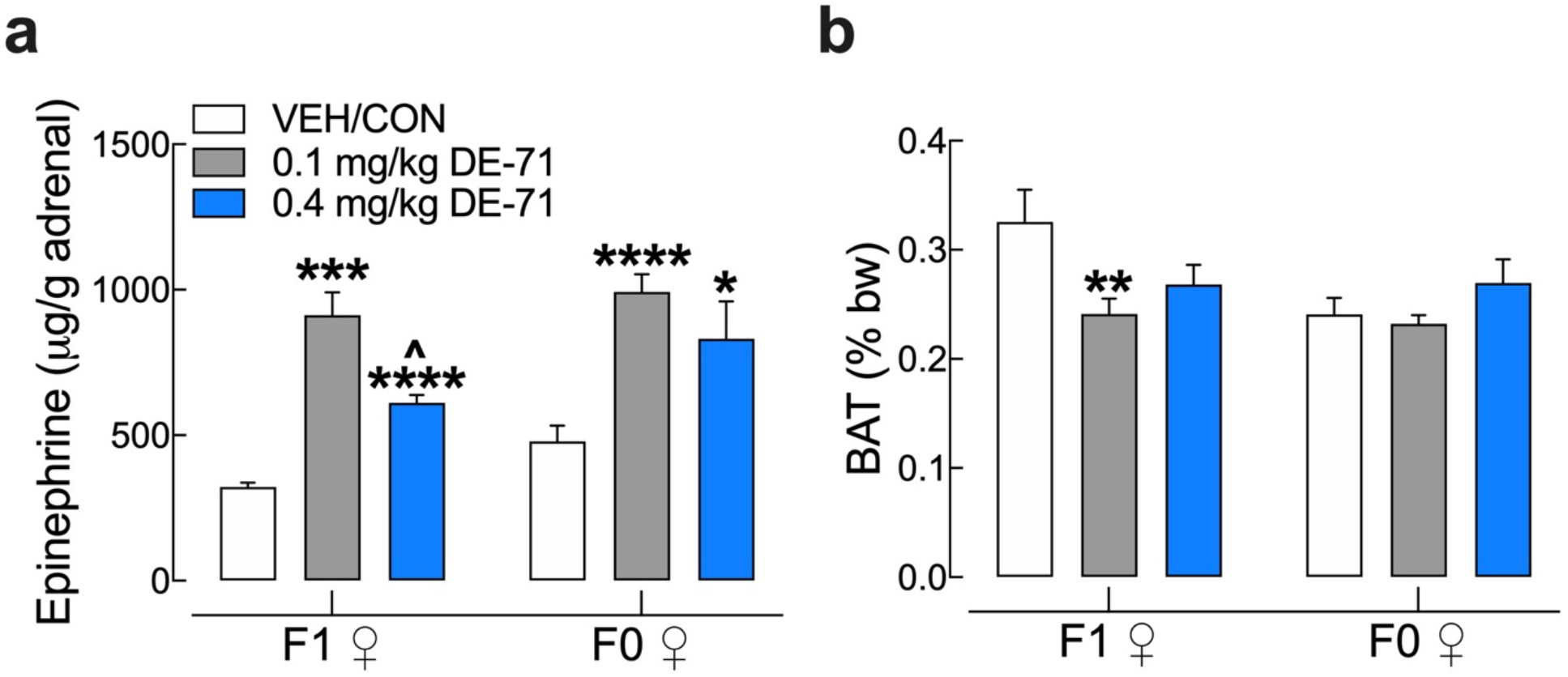
DE-71 exposure increases adrenal epinephrine content in F0 and F1 females and decreases brown adipose tissue (BAT) mass in F1 female mice. **(a)**Epinephrine content measured in adrenal glands harvested at necropsy was elevated in F0 and F1 females exposed to either dose of DE-71. No change in adrenal weight was measured (data not shown). (**b**) Interscapular brown adipose tissue (BAT) collected at necropsy was expressed as a percent of body weight. DE-71 exposure produces reduced BAT mass in F1 at 0.1 mg/kg. No effects were seen in F0. Bars and error bars represent mean±s.e.m. *indicates significantly different from VEH/CON (\**P*<.05, \*\**P*<.01, \*\*\**P*<.001, \*\*\*\**P*<.0001); ^indicates significantly different from 0.1 mg/kg DE-71 (^*P*<0.05). Dunnet’s T3 or Tukey’s *post-hoc* tests were used. Bars and error bars represent mean±s.e.m. n=4-8/group for epinephrine and n=11-32/group for BAT. BAT, brown adipose tissue; F1, female offspring; F0, dams

### DE-71 exposure alters hepatic carbohydrate metabolic enzymatic activity

Elevated glucose levels may be due to hepatic glucose production. Therefore, we tested the hypothesis that PBDEs increase the activity of glutamate dehydrogenase (GDH), a hepatic gluconeogenic enzyme important in normal glucose homeostasis. We found that exposure to DE-71 significantly *reduced* enzymatic activity of GDH in F0 and F1 (**Fig. 8**). DE-71 is significantly more effective at 0.4 mg/kg than at 0.1 mg/kg in both F1 and F0.

**Figure 8.**
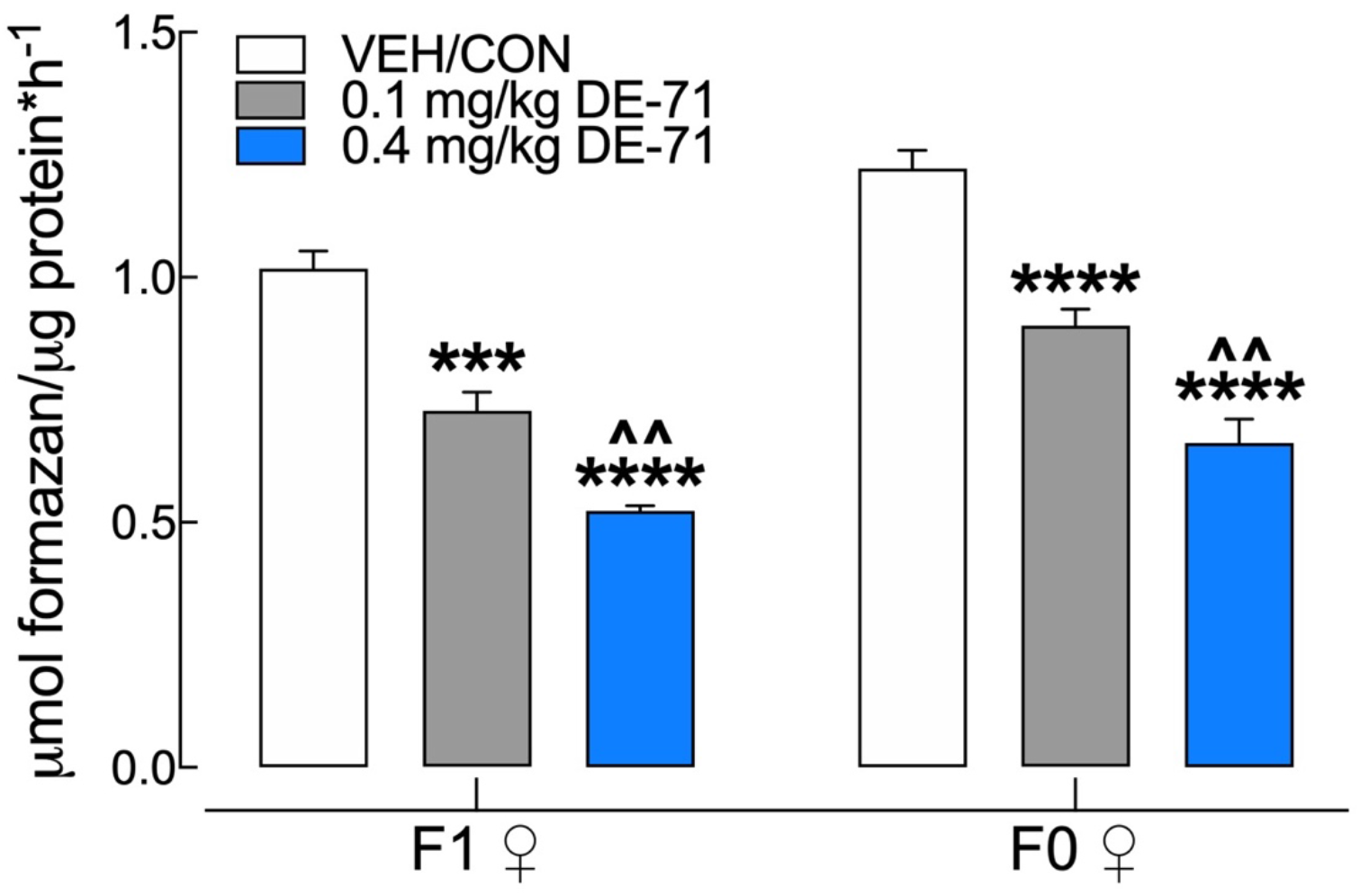
DE-71 exposure reduces hepatic activity of glutamate dehydrogenase (GDH) in F1 and F0 female mice. DE-71 exposure decreased GDH activity in F1 and F0 in a dose-dependent manner, producing greater reduction in the 0.4 mg/kg dose groups. *indicates significantly different from VEH/CON (\*\*\**P*<.001, \*\*\*\**P* <.0001); ^indicates significantly different from 0.1 mg/kg DE-71 (^^*P*<.01). Dunnett’s T3 (F1) or Tukey’s (F0) *post-hoc* tests were used. Bars and error bars represent mean±s.e.m. n=6-8/group. GDH, glutamate dehydrogenase; F1, female offspring; F0, dams

### DE-71 exposure increases hepatic levels of endocannabinoids (EC) and related fatty acid-ethanolamides in exposed F1 but not F0 female mice

F1 exposed to either dose of DE-71 displayed altered liver levels of the endocannabinoid (EC), anandamide (arachidonoylethanolamide, AEA), and related fatty acid-ethanolamides, docosahexaenoyl ethanolamide (DHEA), and n-oleoylethanolamide (OEA), a shorter analogue of AEA, when compared to VEH/CON, with DHEA and OEA being more sensitive to the 0.1 mg/kg dose (**Fig. 9a**). In contrast, no changes were detected for the other primary EC, 2-arachidonoyl-sn-glycerol (2-AG), and related monoacylglycerol, 2-docosahexaenoyl-sn-glycerol (2-DG) (**Fig. 9b**). None of the lipids were altered by DE-71 in exposed F0 females as compared to VEH/CON (**Fig. 9c,d**).

**Figure 9.**
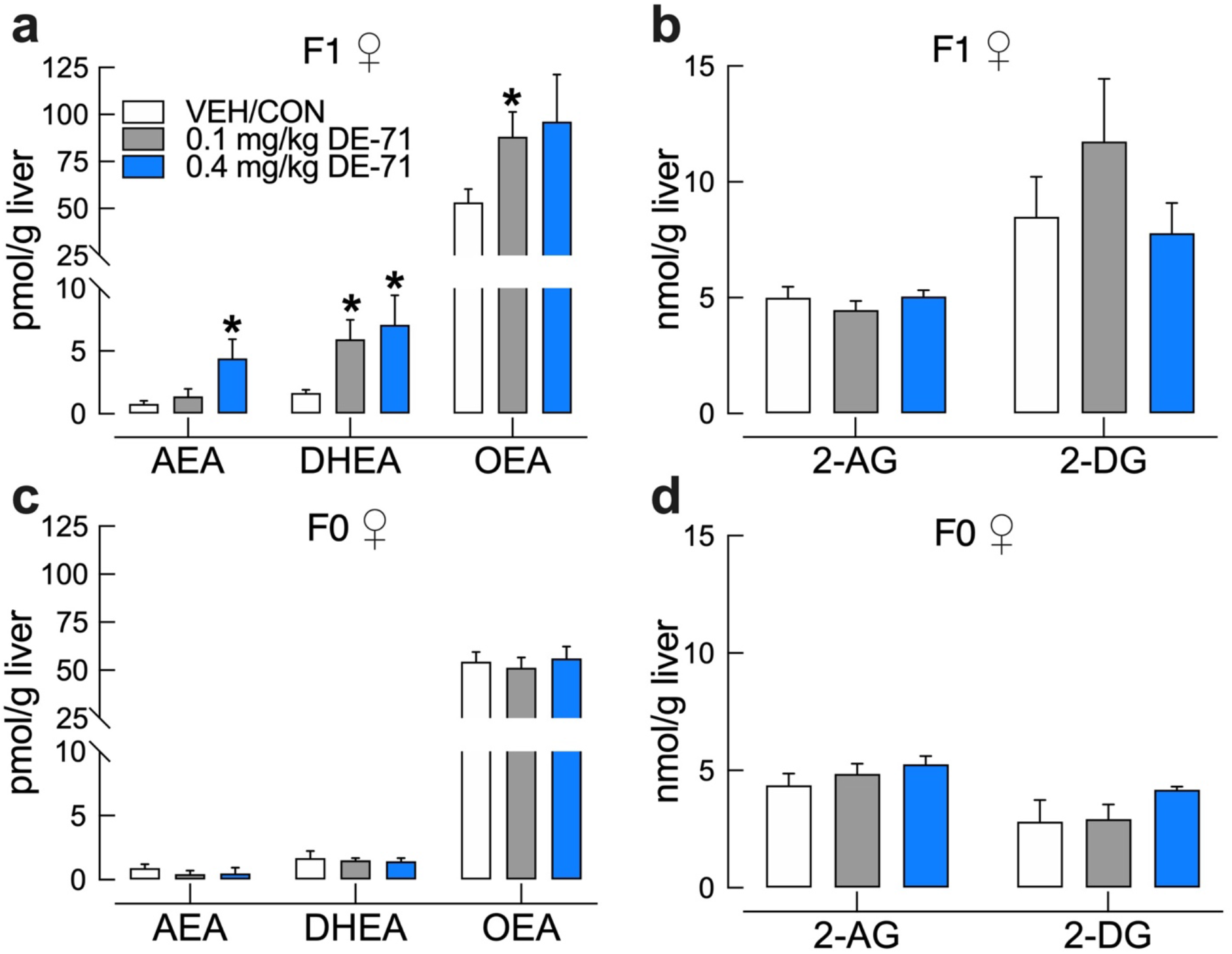
DE-71 exposure increases hepatic levels of endocannabinoid (EC) and related fatty acid-ethanolamides in exposed F1 but not F0 female mice. *Post mortem* liver tissue was analyzed using UPLC/MS/MS. Perinatal (**a,c**) but not adult exposure to DE-71 (**b,d**) produced elevated levels of hepatic ECs and fatty acid-ethanolamides. F1 exposed to 0.1 mg/kg displayed elevated DHEA and OEA (**a**). F1 exposed to 0.4 mg/kg showed elevated AEA (Anandamide), a primary EC, and DHEA. No significant differences were detected after adult exposure to DE-71 in F0 (**b,d**). The other primary EC, 2-AG and 2-DG did not exhibit changes in DE-71 exposed F1 (**b**) or F0 mice (**d**). *indicate significantly different from VEH/CON (\**P*<.05). Bars and error bars represent mean±s.e.m. n=6-16/group for F1 and n=3-5/group for F0. Dunnett’s T3 and Tukey’s *post-hoc* test was used. AEA, arachidonoylethanolamide (Anandamide); DHEA, docosahexanoyl ethanolamide; OEA, n-oleoyl ethanolamide; 2-AG, 2-arachidonoyl-*sn*-glycerol; 2-DG, monoacylglycerol 2-docosahexaenoyl-*sn*-glycerol; F1, female offspring; F0, dams

## Discussion

The diabetogenic effects of PBDEs are not well understood. The objective of this study was twofold. First, we explored whether the environmentally relevant industrial PBDE mixture DE-71, at chronic low doses, could influence *in vivo* and *ex vivo* biomarkers of diabetes. Secondly, we compared the diabetogenic potential of adult versus developmental exposure. Our main finding is that developmental exposure to DE-71 produces significant glucose dysregulation including fasting hyperglycemia, impaired glucose tolerance, insulin insensitivity and delayed glucose clearance/utilization in response to insulin; all symptoms used clinically to diagnose diabetes and validate diabetic animal models^44^.

Our results show a more substantial glucose dyshomeostasis in exposed F1 than in F0 females. FBG was elevated in both 0.1 mg/kg (9 h fast) and 0.4 mg/kg F1 groups (11 h fast) compared to VEH/CON. In addition, F1 females exposed to 0.1 mg/kg showed glucose intolerance after a glucose challenge, such that glycemia reached a greater peak and recovery time to basal levels was extended. This abnormal phenotype was not observed in 0.4 mg/kg when expressing glucose levels as a percent of baseline. A similar but less prominent phenotype was observed in F0. The greater glycemia is unlikely due to enhanced hepatic glucose production from amino acids since DE-71 exposure caused significant reduction in the activity GDH, a key hepatic enzyme regulating amino acid-derived gluconeogenesis^49^. Similar downregulation by DE-71 has been reported for another gluconeogenic enzyme, phosphoenolpyruvate carboxykinase (PEPCK)^50^. Instead, the exaggerated peak glycemia after glucose challenge and the delay in the return to baseline blood glucose may be a result of reduced plasma insulin, reduced insulin sensitivity and/or reduced lean mass (muscle, brain and liver), the principal site of glucose disposal^51^.

Another measure of abnormal glucose homeostasis found in 0.1 mg/kg exposed F1 was insulin insensitivity measured as a significant reduction in the blood glucose reduction rate (K_ITT_) compared to VEH/CON. F1 exposed to either DE-71 dose also showed an increased latency to reach minimum glycemia, which may indicate reduced glucose clearance/utilization. Finally, when expressing glucose levels as a percent of baseline, the glycemic response to insulin challenge showed a slower recovery at 60, 90 and 120 min post injection in F1 exposed to 0.1 and 0.4 mg/kg DE-71 relative to VEH/CON. In support of possible DE-71-induced insulin insensitivity, others have reported an increased glucose:insulin ratio in DE-71 exposed rats^50^. BDE-28 and penta-PBDE have been shown to reduce insulin signaling in adipocytes from insulin-resistant obese humans^36^ and rats^32^, respectively. Moreover, epigenetic/genetic changes in the liver of BDE-47-exposed rat offspring have been associated with insulin signaling and a canonical pathway related to Type 1 diabetes (T1D)^26,28^. Compared to the diabetogenic phenotype of F1 females, adult-exposed dams did not display fasting hyperglycemia nor an abnormal glycemia response to insulin and showed a more subtle glucose intolerance on IPGTT. These results suggest that developmental exposure to environmentally relevant PBDE congeners increases the risk of developing T2D later in life.

Previous experimental studies examining measures of glucose in developmentally exposed female rodents have reported results discordant with ours. No changes in measures of insulin or glucose action were found in female (or male) F1 exposed to a complex BFR mixture found in house dust: DE-71, DE-79, BDE 209 and hexabromocyclododecane (HBCDD) via the mother^52^ possibly explained by a net masking effect of individual congener actions. Using perinatal exposure to BDE-47 Suvorov and colleagues^53^ reported improved glucose uptake on oral GTT in male but not female rat offspring, suggesting sex-specific effects. The differential results of these studies relative to ours may be explained by different perinatal dosing paradigms using dams, such as DE-71 via oral treats vs. BDE-47 via intravenous injection, respectively, and chronic doses of 0.1 and 0.4 mg/kg vs. 6 doses of 0.002 and 0.2 mg/kg bw, respectively. Another study performed in adult females reports no glucose intolerance in *virgin* female rats exposed to BDE-47 in adulthood^54^. In contrast, our exposed adult F0 showed slight but significant glucose intolerance during IPGTT suggesting that they may be more vulnerable to DE-71 under conditions of pregnancy and lactation. Alternatively, the relatively greater susceptibility of exposed F0 females in our study may be due to the combination of PBDE congeners present in DE-71. In adult male rats and mice BDE-47 treatment produces hyperglycemia in one study and diabetic symptoms in others but only when paired in two-hit models^30,31^. Interestingly, our results using DE-71 are similar to those produced by two brominated flame retardants not found in DE-71, BDE-209 (hyperglycemia with reduced insulin)^28^ and Firemaster-550 (apparent glucose intolerance)^27^.

Given the endocrine-disrupting properties of PBDEs^15,55,56^ we hypothesized that DE-71 would disrupt levels of glucoregulatory hormones that serve as diagnostic biomarkers for diabetes. Studies have shown that levels of insulin, glucagon and glucagon-like peptide 1 (GLP-1) are altered in T2D leading to pathological glucose dyshomeostasis^57^. Insulin and glucagon both normalize blood glucose levels under conditions of high and low plasma glucose, respectively. Insulin action facilitates cellular absorption of glucose from the blood whereas glucagon triggers the release of glucose into the blood from liver stores. Reduced plasma insulin in F1 after exposure to 0.1 mg/kg DE-71 may contribute, in part, to the pronounced glucose intolerance seen in this group. Epidemiological studies measuring insulin have reported conflicting reports, either no association with ∑PBDEs (BDE-47 and −153) in Canadian indigenous populations^33^ or a positive association between omental fat levels of BDE-28 and −99 and fasting insulin secretion in obese individuals^36^. Animal studies have failed to show changes in insulin levels with adult exposure to BDE-47^29^ or postnatal exposure to BDE-47^31^ or perinatal exposure to a mixture of BFRs, including PBDEs and HBCDD^52^. In 0.4 mg/kg F1 insulin reduction was less pronounced, and glucagon was decreased, which could explain the less marked glucose intolerance. This coupled with other changes, such as the lack of BAT mass reduction and/or the differential profile of hepatic endocannabinoids at this dose (see below) may help explain dose-related differences in glucose tolerance. In F0 exposed to 0.1 mg/kg DE-71 produced a significant increase in GLP-1 concomitant with glucose intolerance rather than an insulinotropic effect on glycemia. The inhibitory effects of DE-71 on both glucagon and insulin may indicate an altered capacity of ***α***- and ***β***-cell function in the pancreas, respectively. Interestingly, adult exposure to BDE-47 produced reduced gene expression for the rat GLP-1 receptor^29^.

Because of its important role in glucose metabolism and regulation of glucoregulatory hormones we examined if adrenal epinephrine was impacted by DE-71 exposure and found elevated content as compared to VEH/CON especially at 0.1 mg/kg. In F1, this dose also produced fasting hyperglycemia, glucose intolerance, insulin insensitivity and low plasma insulin in exposed F1 (**see Fig. 4**), suggesting the possibility that DE-71 actions on glucose intolerance and insulin reduction and upregulation of the sympathoadrenal system may be related since diabetic animals show elevated adrenal epinephrine^58^. For example, epinephrine stimulates gluconeogenesis, stimulates glycogenolysis either directly or by facilitating glucagon action and inhibits insulin^59^. Interestingly, penta-BDEs enhance adrenergic-stimulated lipolysis in rat adipocytes^32^. Previous reports of *in vitro* exposure to PBDE (and PCBs) show opposite changes in catecholamine (CA) levels in and/or release from cultured chromaffin cells depending on the congener^60,61^. A limitation of our study is that we did not measure plasma levels of epinephrine, although it seems feasible that adrenal and plasma epinephrine may be co-regulated^62^. In support of this, perinatal DE-71 exposure appears to exaggerate adrenal mRNA levels of the major catecholamine synthetic enzyme, tyrosine hydroxylase, in male rats (unpublished observations, Spurgin and Currás-Collazo).

Having found epinephrine alterations, we measured BAT, which is under the trophic influence of ß-adrenergic sympathetic- and insulin-mediated regulation and epinephrine^63,64,65^ Reduced intrascapular BAT mass was uniquely found in 0.1 mg/kg DE-71-exposed F1 as compared to VEH/CON. Lower BAT mass may contribute to fasting hyperglycemia and reduced glucose clearance^48^ and is negatively associated with central obesity and diabetes^63,64,66^. In correspondence, BAT activation of lipolysis and thermogenesis protects against these^67^. Further studies are needed to determine how BAT participates in metabolic health and PBDE-induced glucose dyshomeostasis.

Unbalanced energy homeostasis, including hyperglycemia caused by either diet-induced obesity or T2D diabetes, is associated with elevated concentrations of endocannabinoids (ECs) in liver, visceral fat, serum, pancreas and small intestine epithelium^68,69,70^. These endogenous lipid molecules act via CB_1_Rs in liver to induce glucose production by increasing gluconeogenic genes and promoting fatty acid synthesis^69,71^. Hepatic CB_1_Rs also participate in insulin signaling and glucose uptake^72^. Pharmacological blockade of CB_1_Rs significantly reduces hyperglycemia, improves glucose tolerance and/or insulin sensitivity in obese diabetic Zucker rats or diet-induced obese mice and humans^73,74,75^. In support of a role of ECs in a T2D diabetogenic phenotype, our data demonstrate that mice with the most glucose dysbiosis (DE-71-exposed F1) display unique increases in levels of the EC, AEA, and related fatty acid-ethanolamides, DHEA and OEA, in liver. F1 exposed to 0.1 mg/kg DE-71 showed increased DHEA and OEA but not in AEA. While it is unclear why this group did not show increased AEA levels relative to VEH/CON, it did show increased DHEA, which is likely an agonist for CB1 and CB2 receptors, indicating that the EC system may participate in the diabetogenic phenotype. The upregulated levels of OEA, which is also an agonist at PPARa receptor pathway, may provide protective effects^76^. Future studies using select antagonists of CB1, CB2, or PPAR***α*** receptors could delineate the role of these ECs in DE-71 diabetogenic phenotype. In contrast to AEA, levels of the other primary EC, 2-arachidonoylglycerol (2-AG), in the liver were not different among groups. Indeed, levels of AEA and 2-AG are not always equally impacted by experimental interventions, which may result from differential regulation of individual EC metabolic pathways^77,78,79^. It should be noted that similar EC system changes in pancreas (not measured here) may contribute to the relative deficiency of plasma insulin that we report for DE-71 exposed F1 females^80^.

Congener profiles in the liver were of particular interest since the liver is a key organ regulating glucose homeostasis and xenobiotic metabolism. Mean values for ∑PBDEs concentration were ~15-fold lower in exposed F1 (~200 ng/g lw) than in exposed F0 (~2900 ng/g lw) at the 0.1mg/kg/d dose. Liver levels in exposed F1 are in the range of maximum values reported for human ∑PBDEs concentration in serum (typically lower than liver) of North American populations including Canadian indigenous Inuits and Crees (max 219-402 ng/g lipid wt) and California U.S. women (max ~749.7 ng/g lipid wt)^12,33^. However, they are 5 to 10-fold greater than those in human breast milk and serum measured in different parts of the world in recent years (in ng/g lipid wt): UK 2014 (15)^81^ and China 2016-2017 (2.87)^34^ and California 2009-2012 (41.5)^82^.

The composition of 9 congeners previously detected in the DE-71 lot used has been reported: BDE-99 (45.3%), −47 (33.3%), −100 (8.2%), −153 (3.6%), −154 (3.2%), −85 (2.5%), −139 (0.9%), −138 (0.5%), −28 (0.2%), and −17 (0.1%)^17^. We also detected BDE-66 and −183 but did not detect −138, all of which have been reported in trace amounts in previous studies^41^. In our study, BDE-99 and −153 were the dominant penetrant congeners followed by BDE-47 and −100 in 0.1 mg/kg exposed F0. These were the same primary congeners reported in the DE-71 lot used in this study. Similar results on predominant congeners (BDE-47, −99, −100 and −153) have been found in women sampled recently^12,34,36,81,83^. The fast elimination of BDE-47 from livers of mice^43,84^ and debromination to lower-brominated congeners such as BDE-28 may explain the lower content of BDE-47 in dam liver relative to that of BDE-99 and −153. In contrast to F0, livers of exposed F1 displayed a smaller set of congeners, i.e., only BDE-28/33 and BDE-153. To our knowledge, no previous studies have determined the complete congener profile in female offspring liver after exclusive indirect exposure to DE-71 via maternal transfer. The observation of disproportionately elevated levels of BDE-153 in F1 and F0 was not unexpected since, unlike BDE-47 and BDE-99, it lacks unsubstituted carbons resulting in poor metabolism by the body and allowing for quick absorption and tissue retention, especially in liver^43,85^.

The lack of hepatic BDE-47 and −99 in F1, may be due to the ultra-low doses used and and/or shorter exposure period that their mothers and/or elimination toxicokinetics in F0 dams receiving direct exposure^43,85^. However, transfer of BDE-47 from dams to offspring does occur during both gestation and lactation^86^ and disposed of more slowly in postnatal mouse pups^84^. One possible limitation of our study is that we measured offspring BDE levels in adulthood (~90 d after weaning), and by this time BDE-47 and other DE-71 congeners could have been eliminated, or hydroxylated and, therefore, not detected. Therefore, it is possible that the concentration and range of PBDE congeners were higher/different at a critical developmental period and at the time of the metabolic measurements. Also, we cannot rule out the possibility that impurities in DE-71 arising during production, such as polybrominated biphenyl (PBB) and polybrominated dibenzofurans (PBDFs)^87^, may contribute to diabetogenic effects reported, although evidence for the tetra-PBBs found in DE-71 is lacking.

It is likely that congeners found in F1 liver, BDE-153 and BDE-28, contribute significantly to the pronounced diabetic phenotype seen in DE-71-exposed F1. Adverse diabetogenic symptoms have been associated with both of these PBDE congeners in human serum, breastmilk and children^34,35,36^. BDE-153 is positively associated with diabetes and/or MetS in studies of men and women in China^29,34^, in US^35^ and with fasting hyperglycemia in Canada^33^ but not in other populations^88,89,90^. In particular, Lim and colleagues^35^ reported that the positive association showed an inverted U-shape indicating a significant effect only at low and moderate PBDE exposure which supports the dose-dependent hormesis seen in our study. Moreover, BDE-28 may contribute to insulin-resistant diabetes typical of T2D^36^. Gestational diabetes mellitus (GDM) in healthy US pregnant women sampled from 2013-2015 is also positively associated with high body burdens of BDE-153 and BDE-28^37,38^. Diabetes has increased to pandemic proportions worldwide during the last few decades and we speculate that PBDEs may act as MDCs contributing to this. Our findings may help inform about the potential risks of POP exposure during development contributing to the etiology of diabetes in adulthood.

In conclusion, we demonstrate that chronic low dose perinatal exposure to an environmentally relevant anthropogenic PBDE mixture, DE-71, produces multi-symptom effects related to diabetes: fasting hyperglycemia, glucose intolerance, abnormal sensitivity and glucose clearance after insulin challenge, and increased hepatic endocannabinoid tone, especially after perinatal exposure. DE-71 effects on F0 were more limited indicating that indirect exposure to developing offspring is more detrimental. Other glycemic control effects that may aggravate/accompany DE-71’s diabetogenic-promoting effects occur more generally in exposed F0 and F1, such as reduced insulin and altered glucoregulatory endocrines, exaggerated sympathoadrenal activity and reduced hepatic GDH enzymatic activity. These adverse health effects appear to be associated with maternal transfer of BDE-28 and BDE-153 to F1. Our results indicate that exposed F1 female mice are susceptible to metabolic reprogramming by DE-71 that leads to a diabetogenic phenotype persisting beyond the period of exposure. Our findings warrant additional animal studies that further characterize PBDE-induced diabetic pathophysiology and identify critical developmental windows of greater susceptibility. They also should guide human studies focused on assessing the risk of emerging adult metabolic disease associated with early life PBDE exposure especially in North American populations.

## Methods

### Animals

C57Bl/6N mice were generated using breeders obtained from Charles River (Raleigh, NC) or Taconic Biosciences (Germantown, NY). Mice were group housed 2-4 per cage and maintained in a non-specific pathogen free vivarium on a 12 h light/dark cycle at an ambient temperature (22 ± 3°C) and relative humidity environment (20-70%). Mice were provided rodent chow and municipal tap water *ad libitum*. Procedures on the care and treatment of animals were performed in compliance with the National Institutes of Health *Guide for the Care and Use of Laboratory Animals* and approved by the University of California, Riverside Institutional Animal Care and Use Committee (AUP#20170026).

### Dosing Solutions

Technical pentabromodiphenyl ether (DE-71; Lot no. 1550OI18A; CAS 32534-81-9), a commercial mixture of PBDEs was originally obtained from Great Lakes Chemical Corporation (West Lafayette, IN). DE-71 dosing solutions were prepared to yield two ultra-low doses: 0.1 and 0.4 mg/kg bw per day. These doses contain a similar molar equivalent of BDE-47 used in previous mouse metabolic and neurobehavioral studies^30,91^. Dosing solution (0.05 g/L) was prepared from a stock solution (0.2 g/L) consisting of DE-71 dissolved in corn oil (Mazola) at 65 °C followed by sonication. Vehicle control solution (VEH/CON) contained corn oil without the addition of DE-71. Solutions were made fresh every 6 months.

### DE-71 Exposure and Animal Use

Female virgin mice (PND 30-60) were introduced to cornflakes daily for 1 week. Dams were randomly assigned to one of three exposure groups: corn oil vehicle control (VEH/CON), 0.1 or 0.4 mg/kg bw per day DE-71 (**Fig. 1**). A 10-week dosing regimen was used that included ~4 weeks of pre-conception, plus gestation (3 weeks) and lactation (3 weeks). Offspring were weaned after the lactation period at PND 21 and housed in same-sex groups. This exposure paradigm was chosen to model human-relevant chronic, low-level exposure^34,81,92^ PBDE transfer from mother to infant has been shown to occur during gestation and lactation in humans^22^ and in rodent models^16,86^. Under this regimen, each dam received a daily exposure to DE-71 for an average of 70-80 d and offspring were exposed perinatally for 39 d via mother’s blood and milk. Dams were fed oral treats, (Kellogg’s® cornflakes) infused with dosing solution (2 uL/g bw) daily, except on PND 0 and 1, a method established to ensure ingestion without the stress of oral gavage^86,93^. Consumption was visually confirmed and offspring co-housed with dams were never observed to ingest cornflakes. During the last week of the 4-week pre-conception exposure period dams were mated with an untreated C57Bl/6N male. A 10-week dosing regimen was used as described^86^ to ensure maternal bioaccumulation prior to conception, especially a concern at low doses. In a subset of dams, gestational weight gain and food intake was monitored daily from GD15-18. F0 and F1 female offspring were used *in vivo* and *ex vivo* for analysis of physiological, metabolic and endocrine parameters (**Fig. 1**). Metabolic endpoints for F0 were chosen to be 1-2 weeks post-lactation, at which time dams were ~5 months of age. In order to compare the adult phenotype of F1 these were tested at a comparable age of 4 months. During sacrifice, under terminal isoflurane anesthesia, cardiac blood (0.3-1 mL) was collected and centrifuged at 16,000 × g for 20 min at 4 °C. A cocktail of protease inhibitors and EDTA was added to the plasma fraction and samples stored at −80 °C until immunoassay analyses. Liver, pancreas, spleen, adrenal glands and interscapular BAT were excised and weighed. Plasma, liver and adrenal samples were snap-frozen over dry ice and stored at −80 °C for later analysis of PBDE congener tissue level determination, plasma endocrines, adrenal epinephrine, liver endocannabinoids and enzymatic activity.

### Congener Analysis via Mass Spectrometry

The concentration of 29 BDE congeners in liver samples collected at sacrifice were determined using gas chromatography/mass spectrometry operated in electron capture negative ionization mode (ECNI) as previously described^94^. Samples were analyzed for nine congeners that comprise the DE-71 lot used as described^17^ (BDE-17, 28/33, −47, −85/155, −99, −100, −138, 153, −154) as well as −25, −30, −49, −66, −71, −75, −116, −119, −156, −181, −183, −190, −191, −200/203, −205, −206, −209 that are commonly monitored in the environment. There are only 6 congeners that make up the great majority (96%) of the DE-71 mixture used here: BDE-99 > −47 > −100 > −153 > −154 > −85 that range from 45.0 to 2.54% w/w^17^. Similarly, 5 of these are the most prevalent BDE congeners found in human tissues: BDE-28, −47, −99, −100, and −153^95^ (Center for Disease Control and Prevention, NHANES, 2015). Prior to extraction, two internal standards were spiked into the homogenized tissue, including a monofluorinated BDE congener (F-BDE-69; Accustandard Inc.) and isotopically labeled BDE-209 (^13^C BDE-209; Wellington Laboratories). BDE congeners were extracted from liver tissues (approximately 1 g) using sonication and an aliquot of the extract was used for gravimetric analysis of lipid content. The remaining extract was purified using a Florisil solid phase extraction cartridge. Extracts were spiked with a third standard, isotopically labeled chlorinated diphenyl ether (^13^C CDE-141; Wellington Laboratories) to measure recovery of the internal standards and then analyzed via gas chromatography mass/ spectrometry using ECNI with methane as a reagent gas as previously described. BDE concentrations are expressed as ng/g lipid weight. The method detection limit (MDL) was calculated using 3 times the standard deviation of the laboratory blanks and was equivalent to 0.5 ng/g lw. Recoveries of F-BDE-69 averaged 90.9% ± 6.9% while ^13^C BDE 209 averaged 106% ± 19%.

### Glucose Tolerance test (IPGTT) and Insulin Tolerance Test (ITT)

To perform the intraperitoneal (ip) glucose tolerance test (IPGTT), mice were fasted overnight (ON) for 11h during the metabolically active dark cycle and then injected with glucose (2.0 g/kg bw, i.p.). Tail blood (1uL) was collected and glucose sampled at time 0 (FBG) and at 15, 30, 60, and 120 min post-glucose challenge. Plasma glucose concentrations were measured using a calibrated glucometer (OneTouch Ultra 2, LifeScan Inc.). Seven days following IPGTT, an insulin tolerance test (ITT) was performed on mice fasted ON for 9h and then injected with Humulin R bolus (0.5 U/kg bw, i.p.). Tail blood was collected and glucose sampled at time 0 (FBG), 15, 30, 45, 60, 90 and 120 min post-injection. The area under (AUC) or above the glycemia curve (inverse AUC) was calculated over 0-120 min post injection. To determine *in vivo* insulin sensitivity, the blood glucose reduction rate after insulin administration, K_ITT_, was calculated using the formula 0.693 × t_1/2_^−1^. Half-life (t_1/2_) was calculated from the slope of the blood glucose concentration from 0-15 min post insulin injection, when plasma glucose concentration declines linearly^51^.

### Immunoassays

Plasma collected via cardiac puncture at each necropsy (*ad libitum* fed state) was analyzed for several peptides using commercially available kits according to manufacturer’s instructions. Plasma insulin was measured using commercial ELISA kits (ALPCO and Mercodia). Colorimetric reaction product was read as optical density at 450 nm on a plate reader. The Mercodia assays had a sensitivity of 0.15 mU/L and 1 mU/L in a dynamic range of 0.15-20 mU/L and 3-200 mU/L. The inter-assay coefficient of variability (CV) was 4.9 and 3.4% and intra-assay CV was 5.1 and 4.5%, respectively. The ALPCO insulin ELISA had a sensitivity of 0.019 ng/mL in a dynamic range of 0.025-1.25 ng/mL and inter- and intra-assay CV of 5.7% and 4.5%, respectively. Active glucagon-like peptide-1 (7-36) Amide (GLP-1) was detected by indirect sandwich amide chemiluminescence ELISA (ALPCO) using a luminescence plate reader. This assay had an analytical sensitivity of 0.15 pM in a dynamic range of 0.45-152 pM and inter- and intra-assay CV of 11.6 and 9.5%, respectively. Glucagon was measured by chemiluminescence ELISA (ALPCO), which has a similar assay principle as the GLP-1 assay. This assay has a sensitivity of 41 pg/mL in a dynamic range of 41-10,000 pg/mL and inter- and intra-assay CV of 9.8% and 7.6%, respectively. All kits were specific to rat/mouse hormones. Plasma insulin, active GLP-1 and glucagon were determined by interpolating absorbance or luminosity values using a 4-parameter-logarithmic standard curve.

### Ultra-performance liquid chromatography-tandem mass spectrometry

Hepatic lipids were extracted following a modification of the Folch method^96^. In brief, samples of flash-frozen liver tissue were weighed (10-20 mg) and homogenized in 1 mL of methanol solution containing 500 pmol d_5_-2-arachidonoyl-*sn*-glycerol, 10 pmol d_4_-oleoyloethanolamide, and 1 pmol d_4_-arachidonoylethanolamide as internal standards followed by the addition of 2mL of chloroform, 1 mL of water, and centrifuged at 2000 × g for 15 min at 4 °C. The organic phase was removed and subjected to chloroform extraction. The pooled lower phases were dried under N_2_ gas and resuspended in 0.1 mL methanol:chloroform (9:1). A 1 uL injection was used for analysis of the EC, arachidonoylethanolamide (AEA) and 2-arachidonoyl-sn-glycerol (2-AG), and related fatty acid ethanolamides, docosahexaenoyl ethanolamide (DHEA) and n-oleoylethanolamide (OEA), and monoacylglycerol, 2-docosahexaenoyl-sn-glycerol (2-DG), respectively, was performed via ultra-performance liquid chromatography coupled to tandem mass spectrometry (UPLC/MS/MS) as previously described by us^97^.

### Glutamate dehydrogenase (GDH) activity

GDH is a key enzyme bridging amino acid-to-glucose pathways. GDH activity in crude liver homogenates was assayed using the tetrazolium salt method of Lee and Lardy^98^ with modification for multi-well plates^99^. The 5% (w/v) homogenates of liver (5 mg) were prepared in 0.25 M sucrose solution. The reaction mixture contained 50 μmol/l of substrate (sodium glutamate), 100 μmol/l of phosphate buffer (pH 7.4), 2 μmol/l of iodonitrotetrazolium chloride and 0.1 μmol/l of NAD and distilled water. The reaction, in all the samples, was started by the addition of sample liver homogenate and proceeded for 30 min at 37 °C. The formazan formed was measured as optical density at 545 nm then converted to concentration using a iodonitrotetrazolium formazan (TCI) standard curve fitted with a linear regression model^99^. Protein content was measured using a bicinchoninic acid assay (ThermoFisher). The activity of GDH was expressed as μmol formazan formed per ug protein per h. Samples were run in duplicate.

### Epinephrine Assay

Epinephrine content in adrenal glands was measured using a modification of the trihydroxyindole method. Using this method of catecholamine oxidation at 0 °C, Kelner and colleagues^100^ obtained values for epinephrine content in bovine chromaffin cell lysates that were nearly identical to those measured using HPLC/electrochemical determination. Briefly, 5mg of adrenal tissue homogenates (in 200 μL of 0.05 N perchloric acid) were centrifuged at 15,000 × g at 0 °C for 15 min. Sample supernatant (30 uL) was added to 10% acetic acid (pH 2). Then 60 μL of 0.25% K_2_Fe(CN)_6_ was added to each sample and the mixture was incubated at 0 °C for 20 min. The oxidation reaction was stopped by the addition of 60 μL of a 9 N NaOH solution containing 4mg/ml ascorbic acid (alkaline ascorbate). Fluorescence emission was determined at 520 nm (excitation wavelength at 420 nm) using a fluorescence plate reader. Each sample yielded mean fluorescence intensity units that were converted into epinephrine concentration expressed as μg/g adrenal wet weight using a calibration standards and polynomial curve fitting.

### Statistical Analysis

Data are presented as mean±s.e.m. An unpaired, two-tailed Student’s t-test and Mann-Whitney U test were used for two group comparisons of K_ITT_ and ∑PBDEs congener data, respectively. A one-way analysis of variance (ANOVA) was used to test the main effect of one factor in more than two groups. A Brown-Forsythe ANOVA was used instead if the group variances were significantly different. When normality assumption failed (Shapiro-Wilk test) a non-parametric test was used (Kruskal-Wallis H test). Data for fasting glycemia were analyzed by two-way ANOVA. ITT and IPGTT experiments were analyzed by repeated measures two-way or mixed model ANOVA. The Geisser-Greenhouse correction was used in some cases as noted. ANOVA was followed by *post hoc* testing for multiple group comparisons while reporting multiplicity-adjusted *P* values. Statistical analyses were performed using GraphPad Prism v.8.4.3. Differences were considered significant at *P*<.05. Additional statistical results can be found in **Supplementary Statistical Results online**.

## Acknowledgements

We are grateful to Dr. G. Chompre for suggestions on experimental design and ALPCO for gift of GLP-1 and Glucagon immunoassay kits. We acknowledge Drs. I. Ethell and F. Sladek for gift of C57Bl/6 mice. We acknowledge technical assistance from Dr. P. Deol and J. Evans, A. Dillon (DiPatrizio lab) and M. Denys, G. Lampel, D. Olomi, K. Rabaani, and J. Tran (Currás-Collazo lab). We thank J. Phan for assistance with animal husbandry. This work was supported by UCR Committee on Research grants (M.C.C); UC MEXUS small grant (M.C.C., E.V.K.); Sigma Xi Research Society award (E.V.K.), UCR STEM-HSI (Dept. of Education) award (E.V.K.); MARC U STAR Fellowship (NIH T34 GM062756; G.M.G.); NIH grants DK119498 and DK114978, and TRDRP grant T29KT0232 (N.V.D.) and NIH R01 grant (R01 ES016099, H.M.S.).

## Author Contributions

Conceptualization, M.C.C., E.V.K.; Methodology, M.C.C., E.V.K., N.V.D., D.A.A., H.M.S. J.M.K., B.D.C., G.M.G.; Validation, M.C.C., E.V.K., B.D.C., P.A.P., N.V.D., H.M.S.; Formal Analysis, E.V.K., M.C.C., D.A.A., P.A.P., N.V.D., A.L.P, H.M.S.; Investigation, E.V.K., B.D.C., D.A.A., P.A.P., A.L.P., V.C. J.M.K., G.M.G., A.E.B. and K.R.B.; Writing - Original Draft, E.V.K., M.C.C.; Writing - Review & Editing, M.C.C., E.V.K., H.M.S., N.V.D. A.L.P.; Visualization, E.V.K.; Resources, M.C.C., N.V.D., H.M.S.; Data Curation, E.V.K., M.C.C.; Supervision, M.C.C., E.V.K., , N.V.D., H.M.S. Project Administration, M.C.C., E.V.K., Funding Acquisition, M.C.C., E.V.K., N.V.D., H.M.S. All authors reviewed and approved the final manuscript.

## Additional Information

### Supplementary information

Accompanies this paper.

### Disclaimer

J.M.K. is now a 2nd Lieutenant at the Uniformed Services University, Department of Defense. Her work was performed at the University of California, Riverside before becoming a military officer. However, we want to emphasize that the opinions and assertions expressed herein are those of the authors and do not necessarily reflect the official policy or position of the Uniformed Services University or the Department of Defense.

**Correspondence** and requests for materials should be addressed to M.C.C.

## References

1. Alaee, M., Arias, P., Sjödin, A. & Bergman, A. An overview of commercially used brominated flame retardants, their applications, their use patterns in different countries/regions and possible modes of release. Environ. Int. 29, 683–689 (2003).

2. Trudel, D., Tlustos, C., Von Goetz, N., Scheringer, M. & Hungerbühler, K. PBDE exposure from food in Ireland: optimising data exploitation in probabilistic exposure modelling. J. Expo. Sci. Environ. Epidemiol. 21, 565–575 (2011).

3. Geyer, H. J. et al. Terminal elimination half-lives of the brominated flame retardants TBBPA, HBCD, and lower brominated PBDEs in humans. Organohalogen Compd. 66, 3820–3825 (2004).

4. Petreas, M. et al. High concentrations of polybrominated diphenylethers (PBDEs) in breast adipose tissue of California women. Environ. Int. 37, 190–197 (2011).

5. Pacyniak, E., Roth, M., Hagenbuch, B. & Guo, G. L. Mechanism of polybrominated diphenyl ether uptake into the liver: PBDE congeners are substrates of human hepatic OATP transporters. Toxicol. Sci. 115, 344–353 (2010).

6. Schecter, A. et al. Polybrominated diphenyl ether (PBDE) levels in livers of U.S. human fetuses and newborns. J. Toxicol. Environ. Health A 70, 1–6 (2007).

7. Darnerud, P. O., Eriksen, G. S., Jóhannesson, T., Larsen, P. B. & Viluksela, M. Polybrominated diphenyl ethers: occurrence, dietary exposure, and toxicology. Environ. Health Perspect. 109 Suppl 1, 49–68 (2001).

8. Ohajinwa, C. M. et al. Hydrophobic organic pollutants in soils and dusts at electronic waste recycling sites: occurrence and possible impacts of polybrominated diphenyl ethers. Int. J. Environ. Res. Public Health 16(3), 360 (2019). doi: 10.3390/ijerph16030360

9. Abbasi, G., Li, L. & Breivik, K. Global historical stocks and emissions of PBDEs. Environ. Sci. Technol. 53, 6330–6340 (2019).

10. Klinčić, D., Dvoršćak, M., Jagić, K., Mendaš, G. & Herceg Romanić, S. Levels and distribution of polybrominated diphenyl ethers in humans and environmental compartments: a comprehensive review of the last five years of research. Environ. Sci. Pollut. Res. 27, 5744–5758 (2020).

11. Terry, P. et al. Polybrominated diphenyl ethers (flame retardants) in mother-infant pairs in the Southeastern U.S. Int. J. Environ. Health Res. 27, 205–214 (2017).

12. Hurley, S. et al. Temporal evaluation of polybrominated diphenyl ether (PBDE) serum levels in middle-aged and older California women, 2011-2015. Environ. Sci. Technol. 51, 4697–4704 (2017).

13. Darrow, L. A. et al. Predictors of serum polybrominated diphenyl ether (PBDE) concentrations among children aged 1-5 years. Environ. Sci. Technol. 51, 645–654 (2017).

14. Vuong, A. M. et al. Maternal polybrominated diphenyl ether (PBDE) exposure and thyroid hormones in maternal and cord sera: the HOME study, Cincinnati, USA. Environ. Health Perspect. 123, 1079–1085 (2015).

15. Kodavanti, P. R. S. & Curras-Collazo, M. C. Neuroendocrine actions of organohalogens: thyroid hormones, arginine vasopressin, and neuroplasticity. Front. Neuroendocrinol. 31(4), 479–496 (2010). doi: 10.1016/j.yfrne.2010.06.005.

16. Costa, L. G. & Giordano, G. Developmental neurotoxicity of polybrominated diphenyl ether (PBDE) flame retardants. Neurotoxicol. 28, 1047–1067 (2007).

17. Kodavanti, P. R. S. et al. Developmental exposure to a commercial PBDE mixture, DE-71: neurobehavioral, hormonal, and reproductive effects. Toxicol. Sci. 116, 297–312 (2010).doi: 10.1093/toxsci/kfq105.

18. Chao, H.-R., Wang, S.-L., Lee, W.-J., Wang, Y.-F. & Päpke, O. Levels of polybrominated diphenyl ethers (PBDEs) in breast milk from central Taiwan and their relation to infant birth outcome and maternal menstruation effects. Environ. Int. 33, 239–245 (2007).

19. Barker, D. J. & Osmond, C. Infant mortality, childhood nutrition, and ischaemic heart disease in England and Wales. Lancet 1, 1077–1081 (1986).

20. Zota, A. R. et al. Polybrominated diphenyl ethers (PBDEs) and hydroxylated PBDE metabolites (OH-PBDEs) in maternal and fetal tissues, and associations with fetal cytochrome P450 gene expression. Environ. Int. 112, 269–278 (2018).

21. Lunder, S., Hovander, L., Athanassiadis, I. & Bergman, A. Significantly higher polybrominated diphenyl ether levels in young U.S. children than in their mothers. Environ. Sci. Technol. 44, 5256–5262 (2010).

22. Toms, L.-M. L. et al. Higher accumulation of polybrominated diphenyl ethers in infants than in adults. Environ. Sci. Technol. 42, 7510–7515 (2008).

23. Rose, M. et al. PBDEs in 2-5 year-old children from California and associations with diet and indoor environment. Environ. Sci. Technol. 44, 2648–2653 (2010).

24. Guariguata, L. et al. Global estimates of diabetes prevalence for 2013 and projections for 2035. Diabetes Res. Clin. Pract. 103, 137–149 (2014).

25. De Long, N. E. & Holloway, A. C. Early-life chemical exposures and risk of metabolic syndrome. Diabetes Metab. Syndr. Obes. 10, 101–109 (2017).

26. Suvorov, A. et al. Rat liver epigenome programming by perinatal exposure to 2,2’,4’4’- tetrabromodiphenyl ether. Epigenomics 12, 235–249 (2020).

27. Patisaul, H. B. et al. Accumulation and endocrine disrupting effects of the flame retardant mixture Firemaster® 550 in rats: an exploratory assessment. J. Biochem. Mol. Toxicol. 27, 124–136 (2013).

28. Zhang, Z. et al. Mechanism of BDE209-induced impaired glucose homeostasis based on gene microarray analysis of adult rat liver. Arch. Toxicol. 87, 1557–1567 (2013).

29. Zhang, Z. et al. Environmental exposure to BDE47 is associated with increased diabetes prevalence: Evidence from community-based case-control studies and an animal experiment. Sci. Rep. 6, 27854 (2016).

30. Wang, D. et al. In utero and lactational exposure to BDE-47 promotes obesity development in mouse offspring fed a high-fat diet: impaired lipid metabolism and intestinal dysbiosis. Arch. Toxicol. 92, 1847–1860 (2018).

31. McIntyre, R. L. et al. Polybrominated diphenyl ether congener, BDE-47, impairs insulin sensitivity in mice with liver-specific Pten deficiency. BMC Obes 2, 3 (2015). doi:10.1186/s40608-014-0031-3

32. Hoppe, A. A. & Carey, G. B. Polybrominated diphenyl ethers as endocrine disruptors of adipocyte metabolism. Obesity 15, 2942–2950 (2007).

33. Cordier, S. et al. Association between exposure to persistent organic pollutants and mercury, and glucose metabolism in two Canadian Indigenous populations. Environ. Res. 184, (2020). doi:10.1016/j.envres.2020.109345

34. Han, X. et al. Associations between the exposure to persistent organic pollutants and type 2 diabetes in East China: A case-control study. Chemosphere 241, 125030 (2020). doi:10.1016/j.chemosphere.2019.12503

35. Lim, J.-S., Lee, D.-H. & Jacobs, D. R. Association of brominated flame retardants with diabetes and metabolic syndrome in the U.S. population, 2003-2004. Diabetes Care 31, 1802–1807 (2008).

36. Helaleh, M. et al. Association of polybrominated diphenyl ethers in two fat compartments with increased risk of insulin resistance in obese individuals. Chemosphere 209, 268–276 (2018).

37. Liu, X. et al. A nested case-control study of the association between exposure to polybrominated diphenyl ethers and the risk of gestational diabetes mellitus. Environ. Int. 119, 232–238 (2018).

38. Eslami, B. et al. Association between serum concentrations of persistent organic pollutants and gestational diabetes mellitus in primiparous women. Environmental Research 151, 706–712 (2016).

39. Rahman, M. L. et al. Persistent organic pollutants and gestational diabetes: A multi-center prospective cohort study of healthy US women. Environ. Int. 124, 249–258 (2019).

40. Yessoufou, A. & Moutairou, K. Maternal diabetes in pregnancy: early and long-term outcomes on the offspring and the concept of ‘metabolic memory’. Exp. Diabetes Res. 2011, 1–12 (2011).

41. La Guardia, M. J., Hale, R. C. & Harvey, E. Detailed polybrominated diphenyl ether (PBDE) congener composition of the widely used penta-, octa-, and deca-PBDE technical flame-retardant mixtures. Environ. Sci. Technol. 40, 6247–6254 (2006).

42. Kautzky-Willer, A., Harreiter, J. & Pacini, G. Sex and gender differences in risk, pathophysiology and complications of type 2 diabetes mellitus. Endocr. Rev. 37, 278–316 (2016).

43. Sanders, J. M., Lebetkin, E. H., Chen, L.-J. & Burka, L. T. Disposition of 2,2’,4,4’,5,5’-hexabromodiphenyl ether (BDE153) and its interaction with other polybrominated diphenyl ethers (PBDEs) in rodents. Xenobiotica 36, 824–837 (2006).

44. King, A. J. F. The use of animal models in diabetes research. Br. J. Pharmacol. 166, 877–894 (2012).

45. Laakso, M. Biomarkers for type 2 diabetes. Mol. Metab. 27S, S139–S146 (2019).

46. Jensen, T. L., Kiersgaard, M. K., Sørensen, D. B. & Mikkelsen, L. F. Fasting of mice: a review. Lab. Anim. 47, 225–240 (2013).

47. Amano, A., Tsunoda, M., Aigaki, T., Maruyama, N. & Ishigami, A. Age-related changes of dopamine, noradrenaline and adrenaline in adrenal glands of mice. Geriatr. Gerontol. Int. 13, 490–496 (2013).

48. Lee, P., Greenfield, J. R., Ho, K. K. Y. & Fulham, M. J. A critical appraisal of the prevalence and metabolic significance of brown adipose tissue in adult humans. Am. J. Physiol. Endocrinol. Metab. 299, E601–6 (2010).

49. Karaca, M. et al. Liver glutamate dehydrogenase controls whole-body energy partitioning through amino acid-derived gluconeogenesis and ammonia homeostasis. Diabetes 67, 1949–1961 (2018).

50. Nash, J. T., Szabo, D. T. & Carey, G. B. Polybrominated diphenyl ethers alter hepatic phosphoenolpyruvate carboxykinase enzyme kinetics in male Wistar rats: implications for lipid and glucose metabolism. J. Toxicol. Environ. Health A 76, 142–156 (2013).

51. Ayala, J. E. et al. Standard operating procedures for describing and performing metabolic tests of glucose homeostasis in mice. Dis. Model. Mech. 3, 525–534 (2010).

52. Tung, E. W. Y. et al. Gestational and lactational exposure to an environmentally-relevant mixture of brominated flame retardants: Effects on Neurodevelopment and Metabolism. Birth Defects research 109, 497–512 (2017).

53. Suvorov, A., Battista, M.-C. & Takser, L. Perinatal exposure to low-dose 2,2’,4,4’-tetrabromodiphenyl ether affects growth in rat offspring: what is the role of IGF-1? Toxicol. 260, 126–131 (2009).

54. Krumm, E. A. et al. Organophosphate flame-retardants alter adult mouse homeostasis and gene expression in a sex-dependent manner potentially through interactions with ERα. Toxicol. Sci. 162, 212–224 (2018).

55. Hamers, T. et al. In vitro profiling of the endocrine-disrupting potency of brominated flame retardants. Toxicol. Sci. 92, 157–173 (2006).

56. Stoker, T. E. et al. In vivo and in vitro anti-androgenic effects of DE-71, a commercial polybrominated diphenyl ether (PBDE) mixture. Toxicol. Appl. Pharmacol. 207, 78–88 (2005).

57. Sandoval, D. A. & D’Alessio, D. A. Physiology of proglucagon peptides: role of glucagon and GLP-1 in health and disease. Physiol. Rev. 95, 513–548 (2015).

58. Gallego, M., Setién, R., Izquierdo, M. J., Casis, O. & Casis, E. Diabetes-induced biochemical changes in central and peripheral catecholaminergic systems. Physiol. Res. 52, 735–741 (2003).

59. Exton, J. H. Mechanisms of hormonal regulation of hepatic glucose metabolism. Diabetes. Metab. Rev. 3, 163–183 (1987).

60. Messeri, M. D., Bickmeyer, U., Weinsberg, F. & Wiegand, H. Congener specific effects by polychlorinated biphenyls on catecholamine content and release in chromaffin cells. Arch. Toxicol. 71, 416–421 (1997).

61. Dingemans, M. M. L. et al. Hydroxylation increases the neurotoxic potential of BDE-47 to affect exocytosis and calcium homeostasis in PC12 cells. Environ. Health Perspect. 116, 637–643 (2008).

62. Jobidon, C., Nadeau, A., Tancrède, G., Nguyen, M. H. & Rousseau-Migneron, S. Plasma, adrenal, and heart catecholamines in physically trained normal and diabetic rats. Diabetes 34, 532–535 (1985).

63. Himms-Hagen, J. Brown adipose tissue thermogenesis and obesity. Prog. Lipid Res. 28, 67–115 (1989).

64. Wang, Q. et al. Brown adipose tissue in humans is activated by elevated plasma catecholamines levels and is inversely related to central obesity. PLoS One 6, e21006 (2011).

65. Sharara-Chami, R. I., Joachim, M., Mulcahey, M., Ebert, S. & Majzoub, J. A. Effect of epinephrine deficiency on cold tolerance and on brown adipose tissue. Mol. Cell. Endocrinol. 328, 34–39 (2010).

66. Lowell, B. B. et al. Development of obesity in transgenic mice after genetic ablation of brown adipose tissue. Nature 366, 740–742 (1993).

67. Argyropoulos, G. & Harper, M.-E. Invited Review: Uncoupling proteins and thermoregulation. J. Appl. Physiol. 92, 2187–2198 (2002).

68. Osei-Hyiaman, D. et al. Endocannabinoid activation at hepatic CB1 receptors stimulates fatty acid synthesis and contributes to diet-induced obesity. J. Clin. Invest. 115, 1298–1305 (2005).

69. Matias, I. et al. Dysregulation of peripheral endocannabinoid levels in hyperglycemia and obesity: Effect of high fat diets. Mol. Cell. Endocrinol. 286, S66–78 (2008).

70. Argueta, D. A. & DiPatrizio, N. V. Peripheral endocannabinoid signaling controls hyperphagia in western diet-induced obesity. Physiol. Behav. 171, 32–39 (2017).

71. Chanda, D. et al. Cannabinoid receptor type 1 (CB1R) signaling regulates hepatic gluconeogenesis via induction of endoplasmic reticulum-bound transcription factor cAMP-responsive element-binding protein H (CREBH) in primary hepatocytes. J. Biol. Chem. 286, 27971–27979 (2011).

72. Liu, J. et al. Hepatic cannabinoid receptor-1 mediates diet-induced insulin resistance via inhibition of insulin signaling and clearance in mice. Gastroenterol. 142, 1218–1228.e1 (2012).

73. Janiak, P. et al. Blockade of cannabinoid CB1 receptors improves renal function, metabolic profile, and increased survival of obese Zucker rats. Kidney Int. 72, 1345–1357 (2007).

74. Tam, J. et al. Peripheral CB1 cannabinoid receptor blockade improves cardiometabolic risk in mouse models of obesity. J. Clin. Invest. 120, 2953–2966 (2010).

75. Van Gaal, L. F., Rissanen, A. M., Scheen, A. J., Ziegler, O. & Rossner, S. Effects of the cannabinoid-1 receptor blocker Rimonabant on weight reduction and cardiovascular risk factors in overweight patients: 1-year experience from the RIO-Europe study. ACC Current Journal Review 14, 10 (2005).

76. Brown, I. et al. Cannabinoid receptor-dependent and -independent anti-proliferative effects of omega-3 ethanolamides in androgen receptor-positive and -negative prostate cancer cell lines. Carcinogenesis 31, 1584–1591 (2010).

77. Wang, H. et al. Differential regulation of endocannabinoid synthesis and degradation in the uterus during embryo implantation. Prostaglandins Other Lipid Mediat. 83, 62–74 (2007).

78. Côté, M. et al. Circulating endocannabinoid levels, abdominal adiposity and related cardiometabolic risk factors in obese men. Int. J. Obes. 31, 692–699 (2007).

79. Moradi, H. et al. Circulating endocannabinoids and mortality in hemodialysis patients. Am. J. Nephrol. 51, 86–95 (2020).

80. Gruden, G., Barutta, F., Kunos, G. & Pacher, P. Role of the endocannabinoid system in diabetes and diabetic complications. Br. J. Pharmacol. 173, 1116–1127 (2016).

81. Drage, D. S. et al. Serum measures of hexabromocyclododecane (HBCDD) and polybrominated diphenyl ethers (PBDEs) in reproductive-aged women in the United Kingdom. Environ. Res. 177, 108631 (2019).

82. Guo, W. et al. PBDE levels in breast milk are decreasing in California. Chemosphere 150, 505–513 (2016).

83. Zhang, J. et al. Polybrominated diphenyl ether concentrations in human breast milk specimens worldwide. Epidemiol. 28 Suppl 1, S89–S97 (2017).

84. Staskal, D. F., Diliberto, J. J. & Birnbaum, L. S. Disposition of BDE 47 in developing mice. Toxicol. Sci. 90, 309–316 (2006).

85. Staskal, D. F., Hakk, H., Bauer, D., Diliberto, J. J. & Birnbaum, L. S. Toxicokinetics of polybrominated diphenyl ether congeners 47, 99, 100, and 153 in mice. Toxicol. Sci. 94, 28–37 (2006).

86. Ta, T. A. et al. Bioaccumulation and behavioral effects of 2,2’,4,4’-tetrabromodiphenyl ether (BDE-47) in perinatally exposed mice. Neurotoxicol. Teratol. 33, 393–404 (2011).

87. Hanari, N. et al. Occurrence of polybrominated biphenyls, polybrominated dibenzo-p-dioxins, and polybrominated dibenzofurans as impurities in commercial polybrominated diphenyl ether mixtures. Environ. Sci. Technol. 40, 4400–4405 (2006).

88. Turyk, M. et al. Persistent organic pollutants and biomarkers of diabetes risk in a cohort of Great Lakes sport caught fish consumers. Environ. Res. 140, 335–344 (2015).

89. Airaksinen, R. et al. Association between type 2 diabetes and exposure to persistent organic pollutants. Diabetes Care 34, 1972–1979 (2011).

90. Lee, D.-H. et al. Polychlorinated biphenyls and organochlorine pesticides predict development of type 2 diabetes in the elderly: the prospective investigation of the vasculature in Uppsala seniors (Pivus) study. ISEE Conference Abstracts 2011 (2011).

91. Woods, R. et al. Long-lived epigenetic interactions between perinatal PBDE exposure and Mecp2308 mutation. Hum. Mol. Genet. 21, 2399–2411 (2012).

92. Ongono, J. S. et al. Dietary exposure to brominated flame retardants and risk of type 2 diabetes in the French E3N cohort. Environ. Int. 123, 54–60 (2019).

93. Coburn, C. G., Gillard, E. R. & Currás-Collazo, M. C. Dietary exposure to aroclor 1254 alters central and peripheral vasopressin release in response to dehydration in the rat. Toxicol. Sci. 84, 149–156 (2005). doi: 10.1093/toxsci/kfi046

94. Stapleton, H. M., Dodder, N. G., Offenberg, J. H., Schantz, M. M. & Wise, S. A. Polybrominated Diphenyl Ethers in House Dust and Clothes Dryer Lint. Environ. Sci. Technol. 39, 925–931 (2005).

95. Genuis, S. K., Birkholz, D. & Genuis, S. J. Human excretion of polybrominated diphenyl ether flame retardants: blood, urine, and sweat study. Biomed. Res. Int. 2017, (2017). doi:10.1155/2017/3676089

96. Argueta, D. A., Perez, P. A., Makriyannis, A. & DiPatrizio, N. V. Cannabinoid CB1 receptors inhibit gut-brain satiation signaling in diet-induced obesity. Front. Physiol. 10, 704 (2019).

97. Perez, P. A. & DiPatrizio, N. V. Impact of maternal western diet-induced obesity on offspring mortality and peripheral endocannabinoid system in mice. PLoS One 13(10), e0205021 (2018). doi: 10.1371/journal.pone.0205021.

98. Lee, Y. P. & Lardy, H. A. Influence of thyroid hormones on L-alpha-glycerophosphate dehydrogenases and other dehydrogenases in various organs of the rat. J. Biol. Chem. 240, 1427–1436 (1965).

99. Egido, L. L. D. et al. A spectrophotometric assay for robust viability testing of seed batches using 2,3,5-triphenyl tetrazolium chloride: using *Hordeum vulgare L*. as a model. Front. Plant Sci. 8, 747 (2017). doi: 10.3389/fpls.2017.00747

100. Kelner, K. L., Levine, R. A., Morita, K. & Pollard, H. B. A comparison of trihydroxyindole and HPLC/electrochemical methods for catecholamine measurement in adrenal chromaffin cells. Neurochem. Int. 7, 373–378 (1985).

