## Supplemental Information for "Maternal Transfer of Environmentally Relevant Polybrominated Diphenyl Ethers (PBDEs) Produces a Diabetic Phenotype and Disrupts Glucoregulatory Hormones and Hepatic Endocannabinoids in Adult Mouse Female Offspring"

**Supplementary Table S1.** Dam Gestational Parameters and Litter Outcomes After Chronic Low Dose exposure to DE-71

| | VEH/CON | 0.1 mg/kg/d DE-71 | 0.4 mg/kg/d DE-71 |
|---|---|---|---|
| **Maternal Parameters** | | | |
| n[a] | 7 | 8 | 11 |
| Gestational food intake | | | |
| Absolute (g/day) | 6.83 ± 0.73 | 6.30 ± 0.34 | 6.67 ± 0.31 |
| Relative (g/pup) | 1.20 ± 0.20 | 1.35 ± 0.32 | 1.05 ± 0.15 |
| Gestational weight gain | | | |
| Absolute (g) | 13.98 ± 1.29 | 13.76 ± 0.57 | 13.70 ± 1.23 |
| Relative (g/pup) | 2.39 ± 0.27 | 2.84 ± 0.55 | 2.15 ± 0.37 |
| Relative to initial (%)[b] | 70.50 ± 7.03 | 66.48 ± 4.10 | 65.40 ± 5.46 |
| **Litter Parameters** | | | |
| n | 16 | 19 | 18 |
| Litter size | 5.81 ± 0.57 | 5.79 ± 0.32 | 7.17 ± 0.51 |
| Females/litter | 2.56 ± 0.29 | 3.26 ± 0.33 | 3.61 ± 0.29 |
| Males/litter | 3.25 ± 0.48 | 2.68 ± 0.32 | 3.56 ± 0.40 |
| Secondary sex ratio (M/F) | 0.54 ± 0.04 | 0.46 ± 0.05 | 0.48 ± 0.04 |

[a]indicates number of dams per treatment group
[b]Initial weight was taken at 18 days before pup birth
Data are expressed as mean±s.e.m. Food intake was measured from GD13-GD18. Maternal weight gain was calculated as difference in weight from pre-pregnancy to late gestation (GD 16-18). Relative weight gain was normalized to pre-pregnancy weight (initial) and to total pups in litter (relative). Litter size was measured at PND0 to avoid the confound of infanticide typical of primiparous C57BL/6 dams. The secondary sex ratio was calculated as the proportion of the total pups that are male.

**Supplementary Table S2.** Mass Spectrometric Analysis (GC/ECNIMS) of PBDE congeners in DE-71-exposed mouse dams and their female offspring

| **Compound/Substituents** | **IUPAC Number** | **Dams** | | | **Female Offspring** | | |
|---|---|---|---|---|---|---|---|
| | | VEH/CON | 0.1 mg/kg DE-71 | 0.4 mg/kg DE-71 | VEH/CON | 0.1 mg/kg DE-71 | 0.4 mg/kg DE-71 |
| n | | 4 | 4 | 3 | 4 | 4 | 4 |
| 2,2′,4-tri BDE | BDE 17 | <MDL | <MDL | <MDL | <MDL | <MDL | <MDL |
| 2,3’,4-tri BDE | BDE 25 | <MDL | <MDL | <MDL | <MDL | <MDL | <MDL |
| 2,4,4′-tri BDE, 2′,3,4-tri BDE | BDE 28, 33 | <MDL | 16.9 ± 4.05* | 13.0 ± 2.55[b] | <MDL | 7.04 ± 3.93 | 22.8 ± 8.15* |
| 2,4,6,-tri BDE | BDE 30 | <MDL | <MDL | <MDL | <MDL | <MDL | <MDL |
| 2,2′,4,4′-tetra BDE | BDE 47 | <MDL | 422 ± 110* | 192 ± 151 | <MDL | <MDL | <MDL |
| 2,2′,4,5′-tetra BDE | BDE 49 | <MDL | <MDL | <MDL | <MDL | <MDL | <MDL |
| 2,3′,4,4′-tetra BDE | BDE 66 | <MDL | 17.9 ± 6.18 | 14.6 ± 8.05 | <MDL | <MDL | <MDL |
| 2,3′,4′,6-tetra BDE | BDE 71 | <MDL | <MDL | <MDL | <MDL | <MDL | <MDL |
| 2,4,4′,6-tetra BDE | BDE 75 | <MDL | <MDL | <MDL | <MDL | <MDL | <MDL |
| 2,2′,3,4,4′-penta BDE, 2,2′,4,4′,6,6′-hexa BDE | BDE 85, 155 | <MDL | 119 ± 11.0** | 138 ± 31.2*** | <MDL | <MDL | <MDL |
| 2,2′,4,4′,5-penta BDE | BDE 99 | <MDL | 1305 ± 153** | 743 ± 425 | <MDL | <MDL | <MDL |
| 2,2′,4,4′,6-penta BDE | BDE 100 | <MDL | 426 ± 41.7*** | 267 ± 86.9* | <MDL | <MDL | <MDL |
| 2,3,4,5,6-penta BDE | BDE 116 | <MDL | <MDL | <MDL | <MDL | <MDL | <MDL |
| 2,3’,4,4’,6 –penta BDE | BDE 119 | <MDL | <MDL | <MDL | <MDL | <MDL | <MDL |
| 2,2′,3,4,4′,5′-hexa BDE | BDE 138 | <MDL | <MDL | <MDL | <MDL | <MDL | <MDL |
| 2,2′,4,4′,5,5′-hexa BDE | BDE 153 | <MDL | 652 ± 52.9** | 1655 ± 206****,^^^ | <MDL | 205 ± 27.7* | 1008 ± 102**,^ |
| 2,2′,4,4′,5,6′-hexa BDE | BDE 154 | <MDL | 57.4 ± 5.61* | 68.9 ± 25.6* | <MDL | <MDL | <MDL |
| 2, 3,3′,4,4′,5-hexa BDE | BDE 156 | <MDL | <MDL | <MDL | <MDL | <MDL | <MDL |
| 2,2′, 3,4,4′,5,6-hepta BDE | BDE 181 | <MDL | <MDL | <MDL | <MDL | <MDL | <MDL |
| 2,2′, 3, 4,4′,5′,6-hepta BDE | BDE 183 | <MDL | 39.6 ± 6.94** | 29.5 ± 8.76* | <MDL | <MDL | <MDL |
| 2,3,3′,4,4′,5,6-hepta BDE | BDE 190 | <MDL | <MDL | <MDL | <MDL | <MDL | <MDL |
| 2, 3,3′,4,4′,5′,6-hepta BDE | BDE 191 | <MDL | <MDL | <MDL | <MDL | <MDL | <MDL |
| 2,2’,3,3’,4,5,6,6’-octa BDE, 2,2′,3,4,4′,5,5′,6-octa BDE | BDE 200, 203 | <MDL | <MDL | <MDL | <MDL | <MDL | <MDL |
| 2, 3,3′,4,4′,5,5′,6-octa BDE | BDE 205 | <MDL | <MDL | <MDL | <MDL | <MDL | <MDL |
| 2,2′,3,3′,4,4′,5,5′,6-nona BDE | BDE 206 | <MDL | <MDL | <MDL | <MDL | <MDL | <MDL |
| 2,2′,3,3′,4,4′,5,5′,6,6′-deca BDE | BDE 209 | <MDL | <MDL | <MDL | <MDL | <MDL | <MDL |
| ΣPBDEs | | <MDL | 206 ± 1.30 | 1017 ± 0.37 | <MDL | 2996 ± 1.26 | 2915 ± 1.57 |

Content was measured as ng/g lipid
Values in table are expressed as mean±s.e.m.
ΣPBDEs are reported are geometric mean ± geometric standard deviation
* Indicates a statistically significant n apparent difference relative to VEH/CON by Dunnet’s T3 *post hoc* test following Brown-Forsythe ANOVA or Tukey’s *post hoc* test following One-way ANOVA (P<.05)
^^^ Indicates significantly different relative to 0.1mg/kg group by Tukey’s *post hoc* test following One-way ANOVA (P<.001)
[a] vs VEH/CON 0.4mg/kg P=.06
BDE, brominated diphenyl ether; IUPAC, International Union of Pure and Applied Chemistry; MDL,method detection limit

**Supplementary Table S3.** Chronic Low Dose DE-71 Exposure Has Minimal Effects on Body and Selected Organ Weights

| | **VEH/CON** | **0.1 mg/kg/d DE-71** | **0.4 mg/kg/d DE-71** |
|---|---|---|---|
| **Dams** | | | |
| n | 23 | 26 | 21 |
| Necropsy body weight | 24.19 ± 0.36 | 24.02 ± 0.35 | 25.05 ± 0.59 |
| Liver | | | |
| Absolute | 1.10 ± 0.03 | 1.20 ± 0.04 | 1.11 ± 0.03 |
| Relative | 45.41 ± 0.89 | 49.75 ± 1.46* | 44.33 ± 0.96^ |
| Pancreas | | | |
| Absolute | 0.17 ± 0.01 | 0.18 ± 0.008 | 0.17 ± 0.01 |
| Relative | 6.97 ± 0.34 | 7.60 ± 0.26 | 6.99 ± 0.39 |
| Spleen | | | |
| Absolute | 0.09 ± 0.004 | 0.09 ± 0.002 | 0.09 ± 0.004 |
| Relative | 3.86 ± 0.13 | 3.72 ± 0.11 | 3.74 ± 0.18 |
| **Female Offspring** | | | |
| n | 22 | 30 | 28 |
| Necropsy body weight | 20.80 ± 0.44 | 19.42 ± 0.37* | 21.73 ± 0.39^^^ |
| Liver | | | |
| Absolute | 0.90 ± 0.02 | 0.86 ± 0.02 | 0.99 ± 0.02*,^^^ |
| Relative | 43.41 ± 0.86 | 44.16 ± 0.75 | 45.55 ± 0.73 |
| Pancreas | | | |
| Absolute | 0.14 ± 0.005 | 0.14 ± 0.006 | 0.16 ± 0.006 |
| Relative | 6.95 ± 0.23 | 7.32 ± 0.25 | 7.30 ± 0.31 |
| Spleen | | | |
| Absolute | 0.08 ± 0.002 | 0.07 ± 0.003 | 0.07 ± 0.002 |
| Relative | 3.73 ± 0.11 | 3.75 ± 0.13 | 3.41 ± 0.10 |

Body weights and organ weights (absolute weights) are in grams; organ-weight-to-body-weight ratios (relative weights) are given as mg organ weight/g body weight. Values are reported as mean±s.e.m.
^^^ Indicates significantly different relative to 0.1mg/kg group by Tukey's *post hoc* test following One-way ANOVA ($P<.001$)
^ Indicates significantly different relative to 0.1mg/kg group by Dunnet's T3 *post hoc* test following Brown-Forsythe One-way ANOVA ($P<.05$)
* Indicates an apparent difference relative to VEH/CON by Dunnet's T3 *post hoc* test following Brown-Forsythe One-way ANOVA or Tukey's *post hoc* test following One-way ANOVA ($P<.05$)

**Supplementary Statistical Results:**

**Figure 2:** BDE congener analysis in liver of DE-71-exposed mouse dams and their adult female offspring. (**a**) F1 liver ∑PBDEs, Brown-Forsythe ANOVA: Exposure $F_{(2.0,3.5)}$=91.9, $P$<.001. F0 liver, One-way ANOVA: $F_{(2,8)}$ =17.8, $P$<.001. Liver ∑PBDEs (F1 vs F0), Mann Whitney U Test for F0 mdn= 3034 (2410-3721) and F1 mdn=208 (162-265) groups (U=0, $P$<.05). (**c**) F1 Liver BDE-28/33, One-way ANOVA: Exposure $F_{(2,9)}$=4.9, $P$<.05. F0 Liver BDE-28/33, Brown-Forsythe ANOVA: Exposure $F_{(2,4.6)}$=10.7, $P$<.05. F0 Liver BDE-66, One-way ANOVA: Exposure $F$(2,8)=3.3, n.s. F0 Liver BDE-154, One-way ANOVA: $F_{(2,8)}$=9.3, $P$<.01. F0 Liver BDE-183, $F_{(2,8)}$=12.7, $P$<.01. (**d**) F0 Liver BDE-47, One-way ANOVA: Exposure $F_{(2,8)}$=5.1, $P$<.05. Liver F0 BDE-85/155, One-way ANOVA: Exposure $F_{(2,8)}$=22.9, $P$<.001. Liver F0 BDE-99, One-way ANOVA: Exposure $F_{(2,8)}$=10.0, $P$<.01. Liver F0 BDE-100, One-way ANOVA: Exposure $F_{(2,8)}$=22.3, $P$<.001. Liver F1 BDE-153, Brown-Forsythe ANOVA, $F_{(2,3.4)}$=76.5, $P$<.01. Liver F0 BDE-153, One-way ANOVA: Exposure $F_{(2,8)}$=65.0, $P$<.0001. n=3-4 replicates/group, analyzed in triplicate. ∑PBDE, the sum of concentration of 9 (F0) or 2 (F1) PBDE congeners; F1, female offspring; F0, dams

**Figure 3.** DE-71 exposure produces elevated fasting blood glucose (FBG) and greater glucose intolerance in perinatally exposed female offspring but not than their mothers. (**a**) Offspring FBG, Two-way ANOVA: Time $F_{(1,50)}$=8.7, $P$<.01; Exposure $F_{(2,50)}$=5.0, $P$<.05; Interaction $F_{(2,50)}$=15.2, $P$<.0001. (**b**) Dams FBG, Two-way ANOVA: Time $F_{(1,49)}$=5.2, ns; Exposure $F_{(2,49)}$=0.6, ns; Interaction $F_{(2,49)}$=0.21, ns. n=7-12/group. ns, not significant

**Figure 4.** DE-71 exposure produces greater glucose intolerance after perinatal exposure compared to adult exposure. (**a**) Offspring IPGTT, RM Two-way ANOVA: Time $F_{(4,96)}$=110.7, $P$<.001; Exposure $F_{(2,24)}$=3.9, $P$<.01; Interaction $F_{(8,96)}$=2.7, P<.05. (**d**) Dams IPGTT, Mixed-effects model ANOVA (Geisser-Greenhouse's ε=0.55): Time $F_{(2.2,46.0)}$=90.8, $P$<.0001; Exposure $F_{(2,21)}$=1.5, n.s.; Interaction

$F_{(8,83)}$=2.0, n.s. (**b**) Mean values for integrated area under the IPGTT glucose curve ($AUC_{IPGTTglucose}$). Offspring $AUC_{IPGTTglucose}$, Brown-Forsythe ANOVA: Exposure $F_{(2.0,15.3)}$=4.9, *P*<.05. (**e**) Dams $AUC_{IPGTTglucose}$, Brown-Forsythe ANOVA: Exposure $F_{(2.0,14.9)}$=1.3, ns. (**c**) Offspring Latency to Maximum Glycemia, Brown-Forsythe ANOVA: Exposure $F_{(2,14.8)}$=3.7, *P*<.05. (**f**) Dams Latency to Maximum Glycemia, One-way ANOVA: Exposure $F_{(2,21)}$=1.6, n.s. (**g**) Offspring Percent basal IPGTT, RM Two-way ANOVA: Time $F_{(4,96)}$=83.6, *P*<.0001; Exposure $F_{(2,24)}$=3.2, *P*=.057; Interaction $F_{(8,96)}$=2.9, *P*<.05. (**i**) Dams Percent basal IPGTT, RM Two-way ANOVA: Time $F_{(4,80)}$=69.4, *P*<.0001; Exposure $F_{(2,20)}$=3.4, *P*=.06; Interaction $F_{(8,80)}$=1.6, n.s. (**h**) Offspring $AUC_{IPGTTglucose}$, One-way ANOVA: Exposure $F_{(2,24)}$=3.9, *P*<.05. (j) Dams $AUC_{IPGTTglucose}$, One-way ANOVA: Exposure $F_{(2,20)}$=3.3, *P*=.06. ns, not significant

**Figure 5.** DE-71 exposure causes less glycemia reduction and delayed glucose clearance after insulin challenge in female offspring (F1) but not (F0) their mothers. (**a**) Offspring ITT: RM Two-way ANOVA: Time $F_{(6,156)}$=62.1, *P*<.0001; Exposure $F_{(2,26)}$=7.4, *P*<.01; Interaction $F_{(12,156)}$=3.8, *P*<.0001. (**e**) Dams ITT, Mixed-model ANOVA: Time $F_{(6,165)}$=41.5, P<.0001; Exposure $F_{(2,28)}$=0.3, n.s.; Interaction $F_{(12,165)}$=1.0, n.s. (**b**) Offspring Inverse $AUC_{ITTglucose}$, One-way ANOVA: Exposure $F_{(2,26)}$=4.4, *P*<.05. (**f**) Dams Inverse $AUC_{ITTglucose}$, One-way ANOVA: Exposure $F_{(2,28)}$=0.01, n.s. (**c**) Offspring $K_{ITT}$, One-way ANOVA: $F_{(2,21)}$=3.1, *P*=.06. (**g**) Dams $K_{ITT}$, One-way ANOVA: $F_{(2,20)}$=1.1, n.s. (**d**) Offspring (F1) Latency to Minimum Glycemia, One-way ANOVA: $F_{(2,26)}$=8.5, *P*<.01. (**h**) Dams (F0) Latency to Minimum Glycemia, Brown-Forsythe ANOVA: $F_{(2,16.63)}$=1.7, n.s. (**i**) Offspring (F1) Percent basal ITT, RM Two-way ANOVA: Time $F_{(6,156)}$=71.4, *P*<.0001; Exposure $F_{(2,26)}$=2.4, n.s.; Interaction $F_{(12,156)}$=3.2, *P*<.0001. (**k**) Dams (F0) Percent Basal Glycemia, Mixed effects model ANOVA: Time $F_{(6,167)}$=38.0, *P*<.0001; Exposure $F_{(2,28)}$=0.2, n.s.; Interaction $F_{(12,167)}$=1.0, n.s. (**j**) Offspring $AUC_{ITTglucose}$, One-way ANOVA: Exposure $F_{(2,26)}$=3.1, *P*=.06. (**l**) Dams, $AUC_{ITTglucose}$, One-Way ANOVA: Exposure $F_{(2,27)}$=.21, n.s. Dunnet’s and Tukey’s *post-hoc* tests were used. ns, not significant

**Figure 6.** Endocrine-disrupting effects of DE-71 on glucose regulatory hormones in F0 and F1 female mice. (**a**) Offspring (F1) Plasma Insulin Levels, Kruskal-Wallis test: Exposure H(2)=10.5, $P$<.01. Dam (F0) Insulin Levels, Brown-Forsythe ANOVA: Exposure $F_{(2,41.5)}$=2.1, n.s. (**b**) Offspring Plasma Glucagon Levels, Brown-Forsythe ANOVA: Exposure $F_{(2,10.0)}$=3.7, $P$=.065. Dam (F0) Glucagon Plasma Levels, Brown-Forsythe ANOVA: Exposure $F_{(2,9.9)}$=3.0, $P$=0.1. (**c**) Offspring (F1) Plasma GLP-1 Levels: Exposure, One-way ANOVA: $F_{(2,30)}$=.6, n.s. Dam Plasma GLP-1 Levels, One-way ANOVA: Exposure $F_{(2,23)}$=6.0, $P$<.01 followed by Tukey's *post-hoc* test. Dunnet's, Dunn's and Tukey's *post hoc* tests were used. F1, female offspring; F0, dams; ns, not significant.

**Figure 7.** DE-71 exposure increases adrenal epinephrine content in F0 and F1 females and decreases brown adipose tissue mass in F1 female mice. (**a**) Adrenal Epinephrine Content. Offspring Epinephrine, Brown-Forsythe ANOVA: Exposure $F_{(2,9.3)}$=38.1, $P$<.0001. Dams Epinephrine, One-way ANOVA: Exposure $F_{(2,17)}$=16.3, $P$<.0001. (**b**) Intrascapular brown adipose tissue (BAT) normalized to body weight. Offspring BAT, One-way ANOVA: Exposure $F_{(2,57)}$=4.8, $P$<.05. Dam BAT: Brown-Forsythe ANOVA: Exposure $F_{(2,20.5)}$ =1.5, n.s. Dunnet's T3 or Tukey's *post-hoc* tests were used. F1, female offspring; F0, dams; BAT, brown adipose tissue; ns, not significant.

**Figure 8.** DE-71 exposure reduces hepatic activity of glutamate dehydrogenase (GDH) in F1 and F0 female mice. Offspring (F1) GDH, Brown-Forsythe ANOVA: Exposure $F_{(2,14)}$=59.1, $P$<.0001. Dams (F0) GDH, One-way ANOVA: Exposure $F_{(2,20)}$=46.1, $P$<.0001. F1, female offspring; F0, dams. Dunnett's T3 (F1) or Tukey's (F0) *post-hoc* tests were used.

**Figure 9.** DE-71 exposure increases hepatic levels of endocannabinoid (EC) and related fatty acid-ethanolamides in exposed F1 but not F0 female mice. **(a**) Offspring (F1) AEA, Brown-Forsythe

ANOVA: Exposure $F_{(2,10.6)}$=4.1, $P$<.05. Offspring (F1) DHEA, Welch's ANOVA: Exposure $F_{(2,10.3)}$=6.0, $P$<.05. Offspring OEA, Welch's ANOVA: Exposure $F_{(2,11.6)}$=3.4, $P$=0.069. (**b**) Offspring (F1) 2-AG, One-way ANOVA: Exposure $F_{(2,19)}$=.79, n.s. Offspring 2-DG, Brown-Forsythe ANOVA: Exposure $F_{(2,30)}$=1.15, n.s. (**c**) Dam (F0) AEA, One-way ANOVA: Exposure $F_{(2,9)}$=.64, n.s. Dam (F0) DHEA, One-way ANOVA: Exposure $F_{(2,8)}$=.16, n.s. Dam OEA, One-way ANOVA: Exposure $F_{(2,9)}$=.21, n.s. (**d**) Dam (F0) 2-AG, One-way ANOVA: Exposure $F_{(2,9)}$=.81. Dam (F0) 2-DG, One-way ANOVA: Exposure $F_{(2,9)}$=.95, n.s. Dunnett's T3 and Tukey's *post-hoc* test was used. EC, endocannabinoid; AEA, arachidonoylethanolamide (Anandamide); DHEA, docosahexanoyl ethanolamide; OEA, n-oleoyl ethanolamide; 2-AG, 2-arachidonoyl-*sn*-glycerol; 2-DG, monoacylglycerol; 2-docosahexaenoyl-*sn*-glycerol. ns, not significant
